# Dual targeting of BTK and BCL2 enhances apoptosis in marginal zone lymphoma models: preclinical activity of BGB-16673 and sonrotoclax

**DOI:** 10.1101/2025.11.24.690186

**Authors:** Alberto J. Arribas, Elisa Civanelli, Camilla Scalise, Luciano Cascione, Andrea Rinaldi, Davide Rossi, Emanuele Zucca, Laura Carosella, Stefano Raniolo, Vittorio Limongelli, Francesco Bertoni

**Author notes:** **Corresponding author**: -Prof. Francesco Bertoni, Institute of Oncology Research, via Francesco Chiesa 5, 6500 Bellinzona, Switzerland.

## Abstract

**Background:** BTK and BCL2 represent key therapeutic targets in B-cell lymphomas, including marginal zone lymphoma (MZL). Here, we evaluated the novel BTK degrader BGB-16673, and the second-generation BCL2 inhibitor sonrotoclax in a panel of MZL cell lines, including models with acquired resistance to BTK, BCL2 and PI3K inhibitors, as single agents and in combination.

**Methods:** Cytotoxicity, transcriptomic changes, apoptosis, and protein expression were assessed for BGB-16673 alone and in combination with venetoclax, sonrotoclax, bendamustine, selinexor, lenalidomide, and tazemetostat.

**Results:** BGB-16673 demonstrated single-agent activity in selected MZL cell lines, inducing BTK degradation and transcriptional repression of BCR signaling and MYC targets. Its transcriptional and phenotypic effects overlapped with those of zanubrutinib but also included specific modulation of oxidative phosphorylation genes.

Combination studies revealed that BGB-16673 synergized with other targeted agents, achieving the best results with sonrotoclax, venetoclax, bendamustine, lenalidomide, and rituximab. The mechanism of action of the combination with the BCL2 inhibitors was further evaluated. The combination with sonrotoclax consistently enhanced apoptosis across MZL models, outperforming venetoclax in potency in most cell lines. Sonrotoclax showed over 10-fold higher activity than venetoclax in three MZL lines and demonstrated synergistic interaction with BGB-16673. Mechanistically, the combination leads to BTK degradation and modulation of anti-apoptotic proteins such as BCL2, MCL1, and BCL-XL.

**Conclusion:** Our findings underscore the potential of BGB-16673 as a therapeutic agent, both as a monotherapy and in combination regimens, for treating MZL patients. Additionally, the results also identify the second-generation BCL2 inhibitor sonrotoclax as another drug to be explored in the same patient populations.

## Background

B cell receptor (BCR) signaling and the apoptotic machinery are crucial pathways for the proliferation and survival of B cells. Their pharmacological inhibition with small molecules targeting BTK and BCL2 has proven their clinical efficacy for treating patients affected by lymphoid neoplasms, including marginal zone lymphoma (MZL)^1-6^.

Depending on the country, six BTK inhibitors (ibrutinib, zanubrutinib, acalabrutinib, orelabrutinib, tirabrutinib, and pirtobrutinib) and the first-generation BCL2 inhibitor venetoclax are approved for the treatment of patients with various mature B-cell lymphoid tumors. Notably, data from clinical trials led to approval of ibrutinib ^7^, zanubrutinib ^8^, and orelabrutinib ^9^ specifically for MZL patients. Conversely, clinical data with the first-in-class BCL2 inhibitor venetoclax as a single agent are limited to the patients (overall response (ORR) rate 67%) enrolled in the initial phase 2 study evaluating the drug in the refractory/relapsed (R/R) setting ^6^, albeit additional preclinical and clinical data are available in combination with other drugs ^10-13^.

Unfortunately, resistance to BTK and BCL2 inhibitors occurs, indicating the need for novel approaches and combinations ^3,14-18^.

Small-molecule BTK protein degraders are a novel class of anti-lymphoma agents with promising preclinical and early clinical data ^19-27^. Among these, BGB-16673 is an orally available chimeric degradation-activating compound (CDAC), inducing the cereblon-mediated degradation of BTK ^19,24,28^. It the phase 1 trial for patients with R/R B cell lymphomas, including MZL patients, BGB-16673 has shown a promising activity (ORR of 50%, n.=20) and tolerable safety ^25,28,29^. It is currently in phase 3 trials for patients with R/R chronic lymphocytic leukemia (CLL) (NCT06973187, NCT06970743, NCT06846671).

Sonrotoclax (BGB-11417) is a second-generation BCL2 inhibitor with reported more robust preclinical anti-tumor activity than venetoclax in a few models derived from mantle cell lymphoma (MCL) and diffuse large B-cell lymphoma ^30-32^. In an ongoing phase 1 trial, the preliminary results show an ORR of 67% and 33% of complete responses in the 12 R/R MZL patients ^33^ . Sonrotoclax is currently in Phase 3 trials for patients with R/R MCL or CLL, and in the first-line setting for CLL patients (NCT06073821, NCT06943872, NCT06742996).

Here, we assessed the BTK degrader BGB-16673 in models derived from bona fide MZL, including some in which we have induced secondary resistance to BTK and PI3K inhibitors ^15,16,34-36^, as a single agent and in combination with other molecules, including the two BCL2 inhibitors venetoclax and sonrotoclax, to provide the rationale for future trials in MZL patients.

## Materials and methods

### Cell lines

Cell lines (VL51, SSK41, Karpas1718, HC1, ESKOL, HAIRM, REC1, and MINO) were cultured according to the recommended conditions, as previously described ^37^. All media were supplemented with fetal bovine serum (10% or 20%) and penicillin-streptomycin-neomycin (≈5,000 units penicillin, 5 mg streptomycin, and 10 mg neomycin/mL; Sigma). Human cell line identities were confirmed by short tandem repeat DNA fingerprinting using the Promega GenePrint 10 System kit (B9510). Cells were periodically tested for mycoplasma negativity using the MycoAlert Mycoplasma Detection Kit (Lonza).

### Compounds

BGB-16673, BGB-21704, and sonrotoclax were kindly provided by Beigene. venetoclax, zanubrutinib, bendamustine, selinexor, lenalidomide, and tazemetostat were purchased from Selleckchem (Houston, TX, USA). Rituximab was purchased from Roche (Basel, Switzerland).

### *In vitro* cytotoxic activity

Cells were seeded at the concentration of 20,000/per well in 96-well plates and exposed to BGB-16673 in a 1/10 dilution series ranging from 0.1 nM to 10 μM and assayed by MTT [3-(4,5-dimethylthiazolyl-2)-2, 5-diphenyltetrazoliumbromide], five days after initial treatment, as previously described ^38^. Zanubrutinib, venetoclax, and sonrotoclax were tested as single agents for 5 days followed by MTT assay. BGB-16673 was combined with venetoclax, sonrotoclax, bendamustine, selinexor, lenalidomide, tazemetostat, and rituximab for 5 days followed by MTT assay. Drug doses and dilutions are shown in Supplementary Table S1.

Sensitivity to drug treatments was evaluated by the IC50, calculated based on the 4-parameter logistic regression using the PharmacoGx R package ^39^. The beneficial effect of the combinations versus the single agents was considered according to the Highest Single Agent (HSA) model ^40^, calculated using the SynergyFinder Plus tool ^41^.

### Immunoblotting

Cells were seeded in T25 flasks at a density of 5 million per 10 mL and treated for 24 hours with 0.1% DMSO or different compounds. Subsequently, the protein extraction was performed by incubating all the conditions with the M-PER (Mammalian Protein Extraction Reagent) lysis buffer plus Halt Protease and Phosphatase Inhibitor Cocktail, EDTA-free (100X) for 30 minutes on ice and then centrifuging for 30 minutes at high speed at 4°C. The protein concentration was determined using the BCA protein assay (Pierce Chemical Co, Dallas, TX, USA) and 30 µg of total protein was prepared to load for each sample. Cell lysates were separated on a gradient of 4-20% SDS-polyacrylamide gel by electrophoresis (SDS-PAGE), and then the proteins were transferred on nitrocellulose membranes to be read. Membranes were incubated with primary antibodies overnight, followed by the anti-mouse or anti-rabbit secondary antibodies for 1 hour at room temperature. Enhanced chemiluminescence detection was done following the manufacturer’s instructions (Amersham Life Science, Buckinghamshire, UK). Luminescence was measured by the Fusion Solo S instrument (Witec AG, Sursee, Switzerland). Protein quantification was performed using the Fusion Solo S instrument (Witec AG). Equal loading of samples was confirmed by probing for GAPDH or vinculin as a housekeeping gene. The antibodies used for the experiment were: anti-BTK (D3H5, CST-8547; Cell Signaling, Danvers, MA, USA), anti-p-BTK (Tyr223) (D9T6H, CST-87141; Cell Signaling), anti-GAPDH (FF26A; eBioscience, San Diego, CA, USA), anti-BLC2 (D55G8, CST-4223; Cell Signaling), anti-MCL1 (D35A5, CST-5453; Cell Signaling), anti-BCL-XL (54H6, CST-2764; Cell Signaling), anti-vinculin (E1E9V, CST-13901; Cell Signaling).

### Apoptosis induction

Cells were seeded at a 2 million per 10 mL density and subsequently treated with the compounds for 24 hours. Cells were stained with Annexin V-FITC (ThermoFisher Scientific, Waltham, MA, USA), washed, and then stained with Propidium Iodide (PI). Acquisitions were carried out with a FACSCanto II instrument (BD Biosciences, Allschwil, Switzerland), and data were analyzed using FlowJo software (TreeStar Inc., Ashland, OR USA). For the cell cycle, cells were fixed with 70% cold ethanol before staining with PI and RNase (Sigma Aldrich, Buchs, Switzerland) treatment. Acquisitions were carried out with a FACSCanto II instrument (BD Biosciences, Allschwil, Switzerland), and data were analyzed using FlowJo software (TreeStar Inc., Ashland, OR, USA). Transcriptome analysis RNA was extracted and processed for RNA sequencing (RNA-Seq; stranded, single-ended 120 bp-long sequencing reads) using the NEBNext Ultra Directional RNA Library Prep Kit for Illumina (New England BioLabs Inc., Ipswich, MA, USA) on a NextSeq 2000 (Illumina, San Diego, CA, USA).

### Data mining

For RNA-Seq data, we evaluated the reads quality with FastQC (v0.11.5) ^42^ and removed low-quality reads/bases and adaptor sequences using Trimmomatic (v0.35) ^43^. The trimmed-high-quality sequencing reads were aligned using STAR ^44^ to the reference genome (HG38). The HTSeq-count software package ^45^ was then used to quantify gene expression with GENCODE v22 as gene annotation. Expression values are provided in a tab-delimited format. We sub-setted the data to genes with a counts-per-million value greater than five in at least one sample. The data were normalized using the ‘TMM’ method from the edgeR package ^46^ and transformed to log2 counts-per-million using the edgeR function ‘cpm.’ The differential gene expression of each comparison of interest was computed using moderated t-test based on TMM-normalization and voom transformation ^47^. Functional analysis was performed using GSEA (Gene Set Enrichment Analysis) with the MSigDB (Molecular Signatures Database) C2-C7 gene sets (18, 19) ^48^, and SignatureDB database ^49^. Statistical tests were performed using the R environment (R Studio console; RStudio, Boston, MA, USA).

### Molecular Docking

The 3D structure of BGB-16673 was generated by converting its Simplified Molecular Input Line Entry System (SMILES) representation into a PDB file using OpenBabel ^50^. The protonation state of the molecules was predicted using the MolGpKa software

^51^. Hydrogen atoms were added using UCSF Chimera ^52^. Rigid-body molecular docking was performed using AutoDock 4.2 ^53^. Ligand and receptor files in PDBQT format were generated via AutoDockTools ^53^, including the calculation of Gasteiger charges and the assignment of rotatable bonds. A 3D potential grid was generated with AutoGrid, with a grid box size of 84 Å × 58 Å × 72 Å and a spacing of 0.375 Å, centered on the coordinates of the crystallographic ligand L0Z (x = 9.192, y = –11.554, z = –6.084).

Docking was performed using the Lamarckian Genetic Algorithm (LGA) with a population size of 150 individuals, a maximum of 2,500,000 energy evaluations, and 500 independent docking runs. The resulting poses were clustered using a root mean square deviation (RMSD) threshold of 2.0 Å. Visual inspection was performed using UCSF Chimera.

## Results

### The BTK-degrader BGB-16673 has single-agent activity in individual MZL

The effect of the BTK-degrader BGB-16673 on cell viability was assessed after five days of exposure in six MZL cell lines, plus two BTK inhibitor-sensitive MCL cell lines, which we included as positive controls (Supplementary Figure 1A; Supplementary Table S2). The SSK41 and Karpas1718 were highly sensitive, with IC50 values of 0.27 and 1.89 nM, respectively (Figure 1). A minor response was observed in the MZL VL51 cell line. No effect was observed in the remaining three MZL models (HC1, HAIRM, and ESKOL) with concentrations of BGB-16673 up to 10 μM. In the two sensitive MZL cell lines, the effect of BGB-16673 overlapped with what was obtained with zanubrutinib (Figure 1B, Supplementary Figure 1B). BGB-16673 was also very active in the two MCL cell lines: REC1 (IC50 0.001 nM) and MINO (0.03 nM) (Figure 1A; Supplementary Table S2).

**Figure 1.**
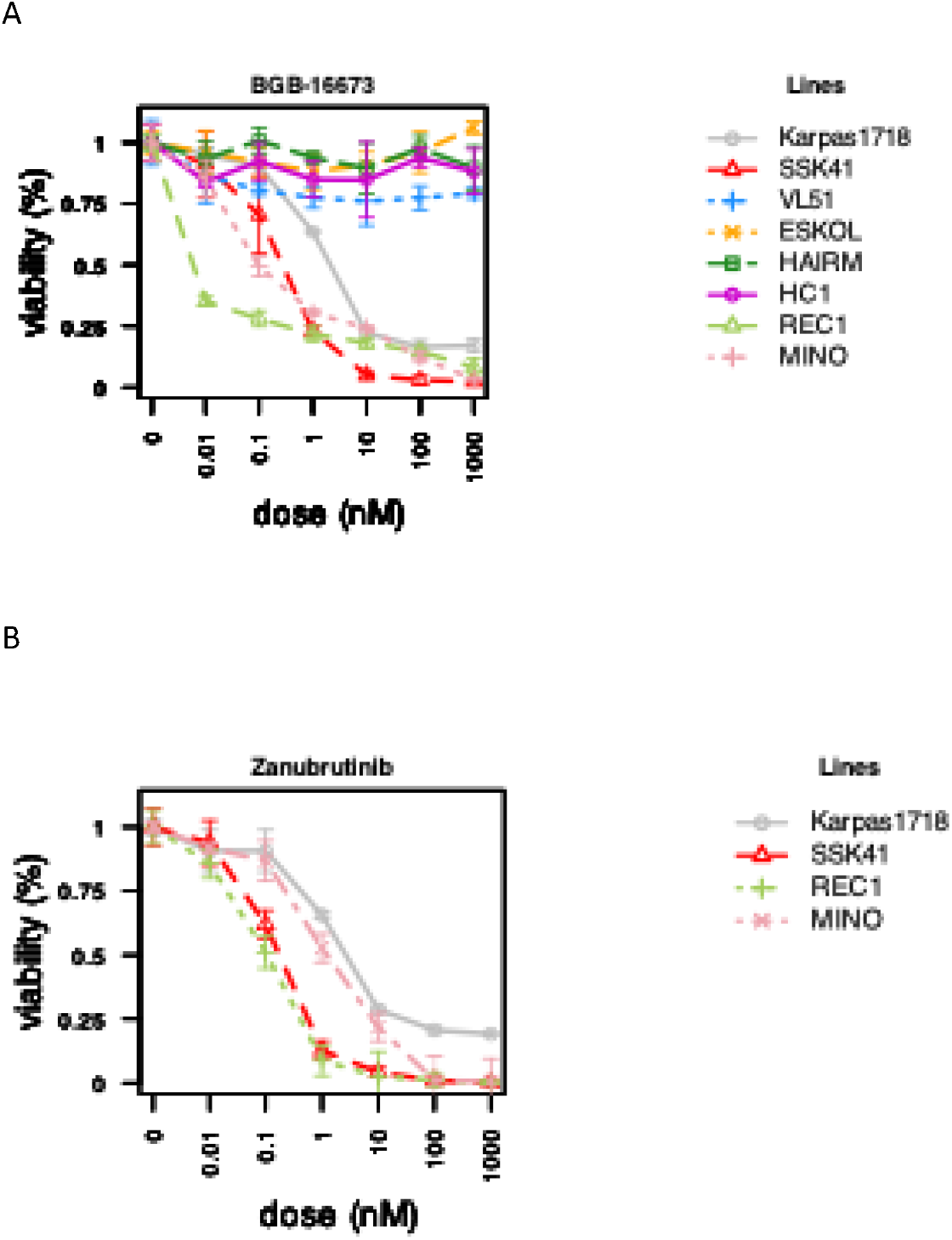
Effect of five days of exposure to the BTK-degrader BGB-16673 (A) or to the BTK inhibitor zanubrutinib (B) in cell lines derived from MZL (VL51, SSK41, Karpas1718, HC1, HAIRM, ESKOL) and from MCL (REC1, MINO).

Both zanubrutinib and BGB-16673 down-regulated the levels of p-BTK(Tyr223), but only the latter down-regulated the total BTK protein levels in all six MZL cell lines (Supplementary Figure 2). The effect followed different kinetics, but it appeared that overall BTK expression is not a major determinant of response (Supplementary Figure 2). However, upon exposure to BGB-16673, resistant lines retained significantly higher levels of phosphorylated BTK (p-BTK), either itself (p=0.044) or p-BTK/BTK ratio (p=0.042), compared to sensitive ones (Supplementary Figure 2H), suggesting that resistant cells maintain active BTK signaling despite drug exposure.

A lower response was observed using BGB-21704, a compound similar to BGB-16673 but with no BTK-related activity, compared to the BTK degrader (Supplementary Figure 3).

Reduced activity was observed in a panel of cell lines derived from the VL51, Karpas1718, and SSK41, which exhibited secondary resistance to BTK, PI3K, and BCL2 inhibitors (Supplementary Figure 4).

### The BTK-degrader BGB-16673 and the BTK inhibitor zanubrutinib induce similar transcriptome changes in MZL cell lines

To compare the mechanism of action between the BTK degrader and the BTK inhibitor, we performed RNA-Seq on RNA extracted from the Karpas1718 cell line exposed to BGB-16673 (5 nM), zanubrutinib (5 nM), or, as a control, to DMSO for 8, 12, and 24 hours (Figure 2, Supplementary Table S3, Supplementary Figure 5). As shown in Figure 2B, the effect of the two compounds on the cell line was very similar (r=0.58, P < 0.001), with very few transcripts showing higher or lower changes with one or the other compound (Figure 2B). Both agents downregulated important gene setsκ, such as NF-κB and B cell receptor signaling, and showed overlapping signatures with other BTK-targeting compounds (Figure 3; Supplementary Table S3). Still of potential interest for combination studies, only BGB-16673 appeared to downregulate transcripts involved in oxidative phosphorylation and ribosome machinery (Figure 3).

**Figure 2.**
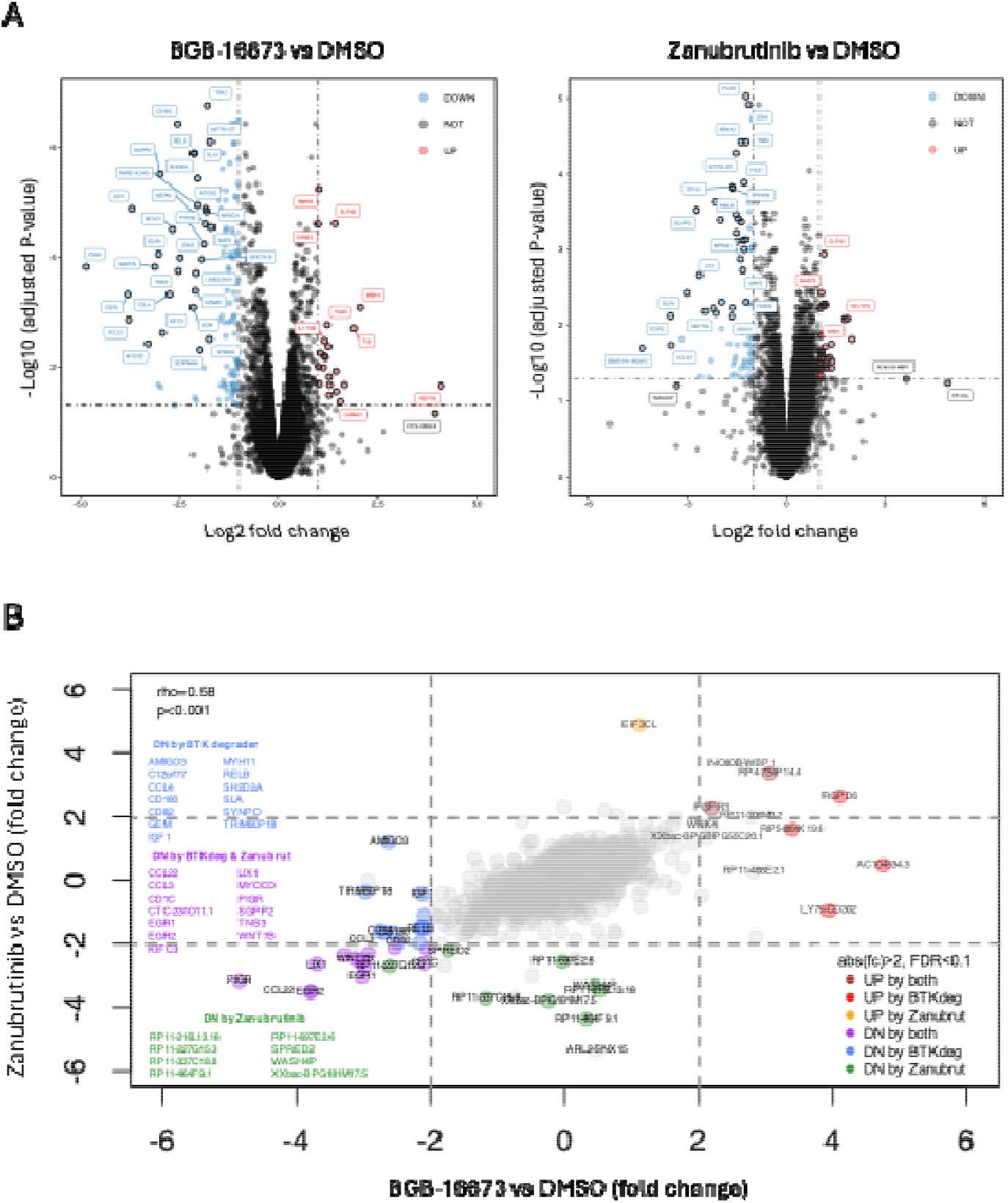
Transcriptome changes in the Karpas1718 cell line after exposure to BGB-16673 (5 nM), zanubrutinib (5 nM), or, as a control, to DMSO for 8, 12, and 24 hours. (A) Volcano plots represent the log2 fold-change in the X-axis and -log10 adjusted P-value in the Y-axis for BGB-16673 (left) or zanubrutinib (right) compared to DMSO. Statistically significant upregulated genes are in red, and downregulated in blue. (B) Correlation plot of the fold-change of transcripts after BGB-16673 (X-axis) or zanubrutinib (Y-axis) versus DMSO.

**Figure 3.**
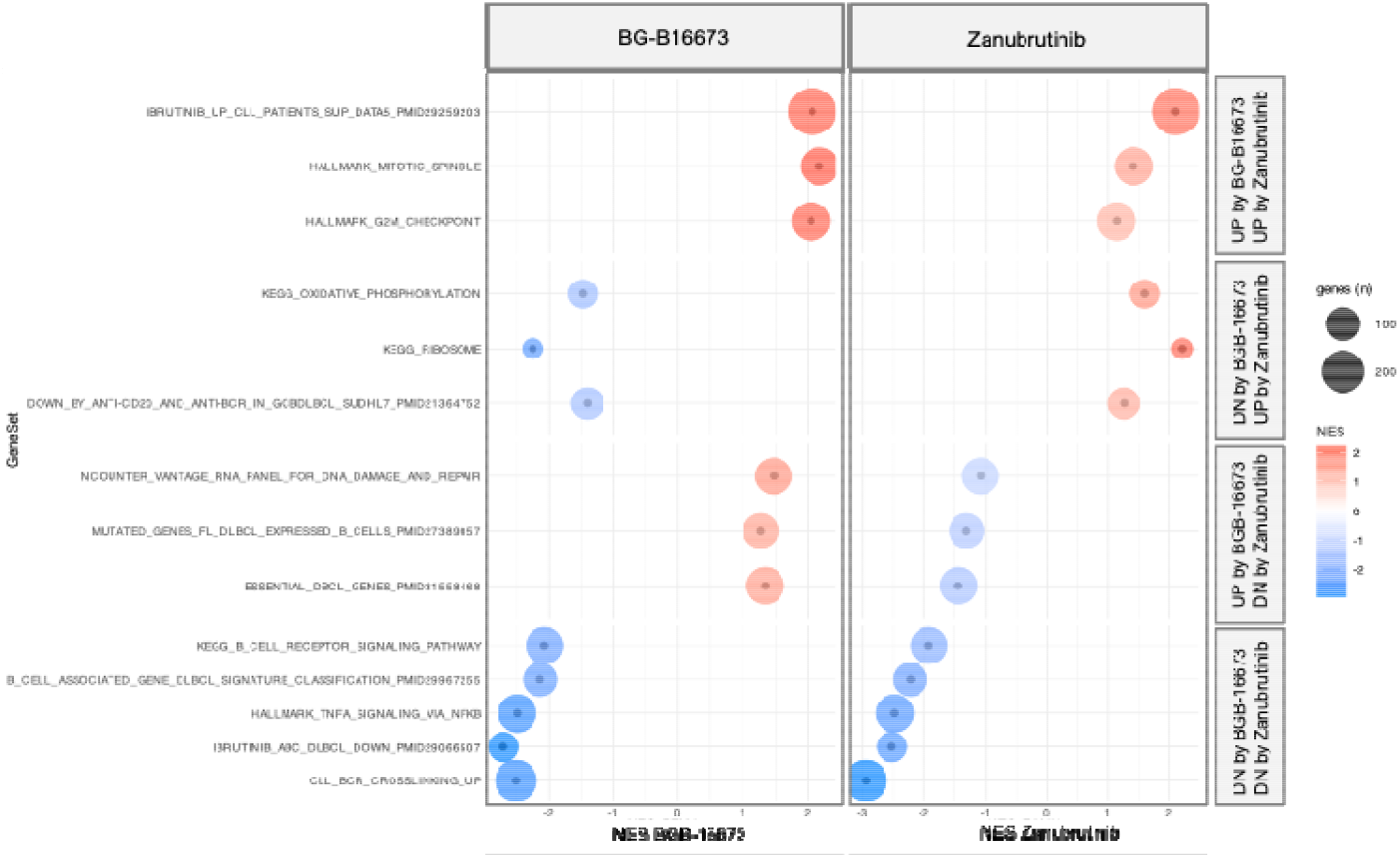
Gene sets modulated by BGB-16673 and zanubrutinib by Gene Set Enrichment Analyses (GSEA). Dot plots representing the changes in pathways upon exposure to BGB-16673 (left) or zanubrutinib (right) in Karpas1718 cells, including the gene count (number of DEG in pathway), and the NES, normalized enrichment score. All gene sets are statistically significant (P>0.05, adjusted P<0.25, |NES|>1).

### The BTK-degrader BGB-16673 synergizes with other targeted agents in MZL cell lines

To explore the potential benefits of BGB-16673-based combinations, we exposed four MZL cell lines to increasing concentrations of the BTK degrader and a series of agents used in clinical settings or in clinical development for a period of five days. The BTK degrader was combined with the two BCL2 inhibitors venetoclax and sonrotoclax, the cereblon E3 ligase modulator lenalidomide, the XPO1 inhibitor selinexor, the EZH2 inhibitor tazemetostat, the DNA alkylating agent bendamustine, and the CD20 targeting monoclonal antibody rituximab (Table 1; Figure 4, Supplementary Figure 6). Considering additivity and synergism, most combinations were beneficial across the cell lines. The best results were obtained when we combined BGB-16673 with the BCL2 inhibitors venetoclax or sonrotoclax, with bendamustine, lenalidomide, and rituximab.

**Table 1.**
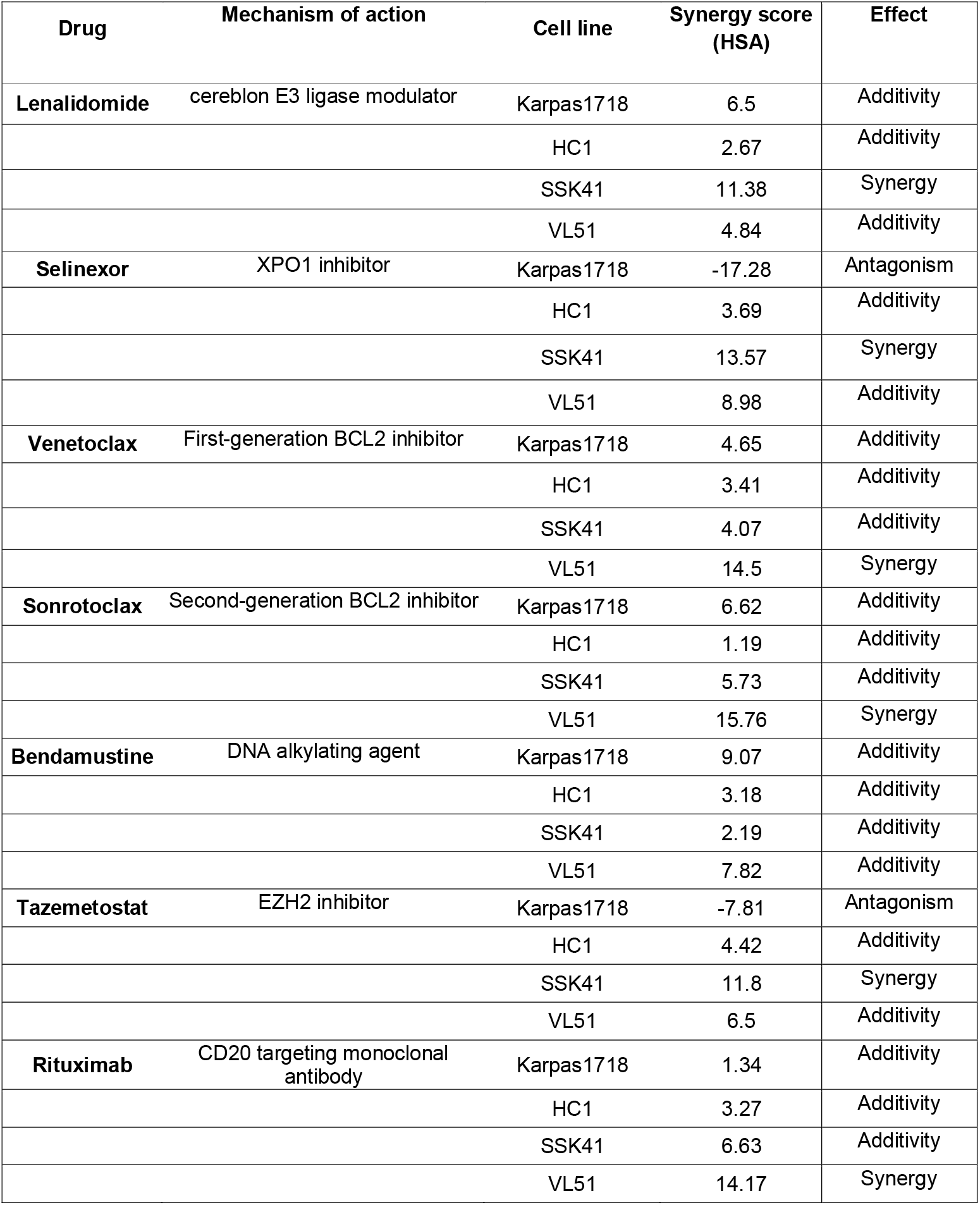
BGB-16673-based combinations in four MZL lymphoma cell lines. Results were obtained via MTT after five days of exposure. HSA, highest single agent.

**Figure 4.**
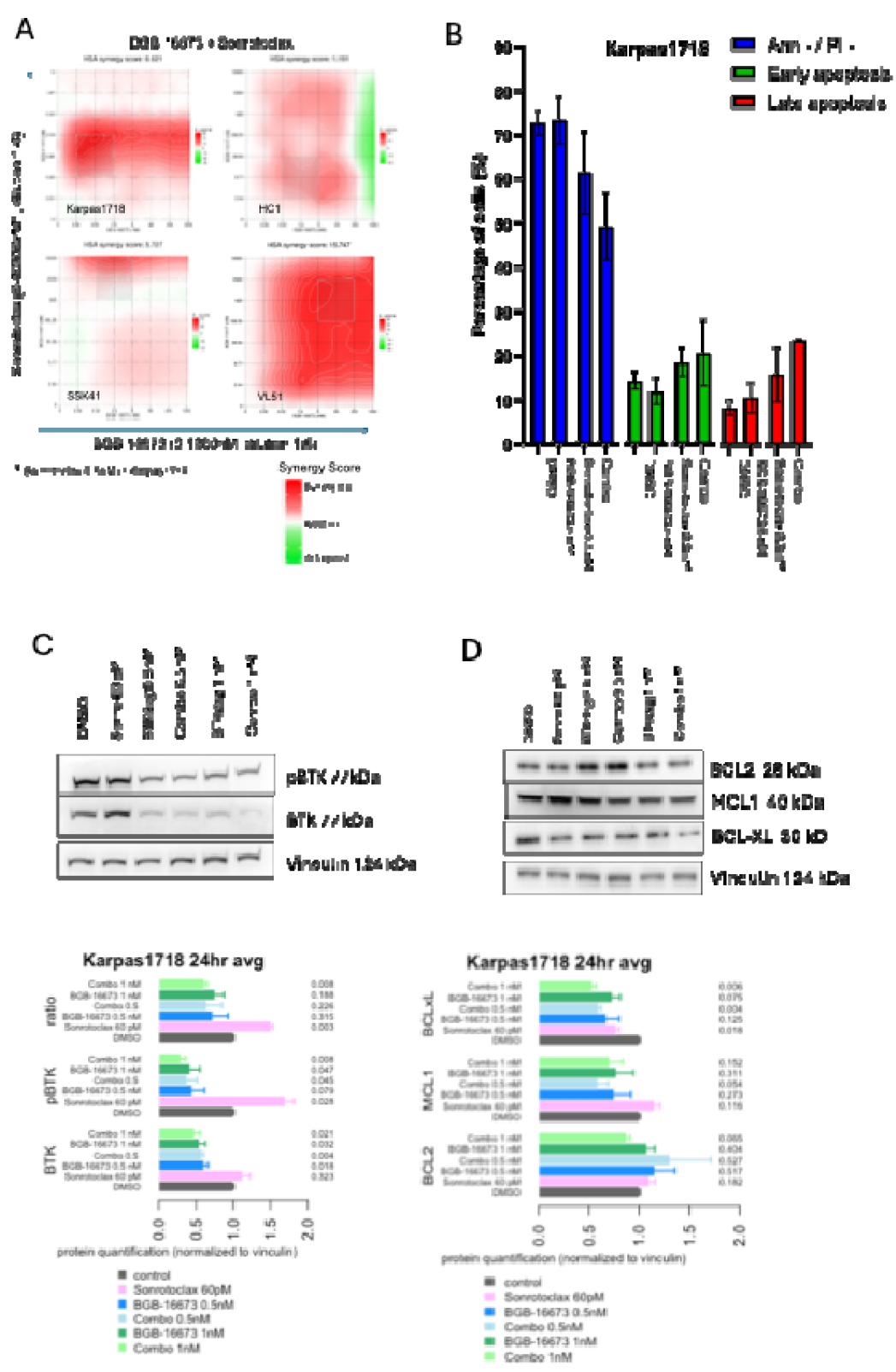
Effect of the combination of BGB-16673 with the BCL2 inhibitor sonrotoclax. Karpas1718, HC1, SSK41 and VL51 MZL cell lines were exposed for five days to increasing concentrations of the BTK degrader and Sonrotoclax, followed by an MTT assay. (A) The benefit of the combination was assessed according to the Highest Single Agent (HSA) algorithm in the Synergy Finder online tool (https://synergyfinder.aittokallio.group/). Red for synergistic, green for antagonistic. (B) Apoptosis induction by BGB-16673 as a single agent and in combination with sonrotoclax in Karpas1718 cells. The bar plots show the mean of two biological replicates after 24 hours of treatment. Error bars represent the standard deviation of the mean. Effect of the BTK-degrader BGB-16673 and the BCL2 inhibitor sonrotoclax on the protein levels of BTK and pBTK (C), and of BCL2, MCL1 and BCL-xL (D) in Karpas1718 cells (24hr of exposure). Representative membranes of two independent experiments. The bar plots show the mean of the protein quantification in the two experiments. Error bars correspond to the standard deviation of the mean, and numbers represent the p-value from a t-test comparing each treatment to the control (DMSO, grey bar).

When we assessed apoptosis induction after 24 hours of exposure to BGB-16673 as a single agent or in combination with venetoclax, sonrotoclax, or bendamustine, we observed higher apoptosis with the combinations than with the single agents (Figure 4, Supplementary Figure 7). Also considering the clinical potential, we then focused on combining BGB-16673 and sonrotoclax to better understand the mechanisms underlying the observed synergy. We investigated the changes in protein levels of p-BTK(Tyr223)/BTK, BCL2, MCL1, and BCL-XL in VL51 and Karpas1718 MZL models. The exposure to BGB-16673 decreased total BTK and p-BTK(Tyr223) levels in both cell lines. The effect was maintained in cells exposed to both BGB-16673 and sonrotoclax (Figure 4, Supplementary Figure 8).

Both Karpas1718 and VL51 cells presented an up-regulation of BCL2 after exposure to the lower concentration of the BTK degrader (0.5 nM) as a single agent and when combined with sonrotoclax. No changes were observed when cells were exposed to 1nM of BGB-16673 as a single or in combination (Figure 4, Supplementary Figure 9).

Sonrotoclax as a single agent increased levels of MCL1 in VL51 cells, with a slight increase in Karpas1718 cells. On the other hand, the BTK degrader BGB16673 repressed MCL1 in Karpas1718 when given as a single agent and in combination with sonrotoclax. In VL51 cells, the BTK degrader up-regulated MCL1 at 1 µM with no effect at lower doses. Finally, the BTK degrader downmodulated BCL-XL in Karpas1718 but not in VL51 (Figure 4, Supplementary Figure 9).

### The BCL2 inhibitor sonrotoclax is more active than venetoclax in MZL models

The combination experiments also allowed a direct comparison between the first- and second-generation BCL2 inhibitors in four cell lines: sonrotoclax was over ten times more active than venetoclax in three cell lines (HC1, HAIRM and Karpas1718), showed improved activity at low doses in models (VL51 and ESKOL, and equally active in the remaining cell line (SSK41) (Figure 5; Supplementary Table S4).

**Figure 5.**
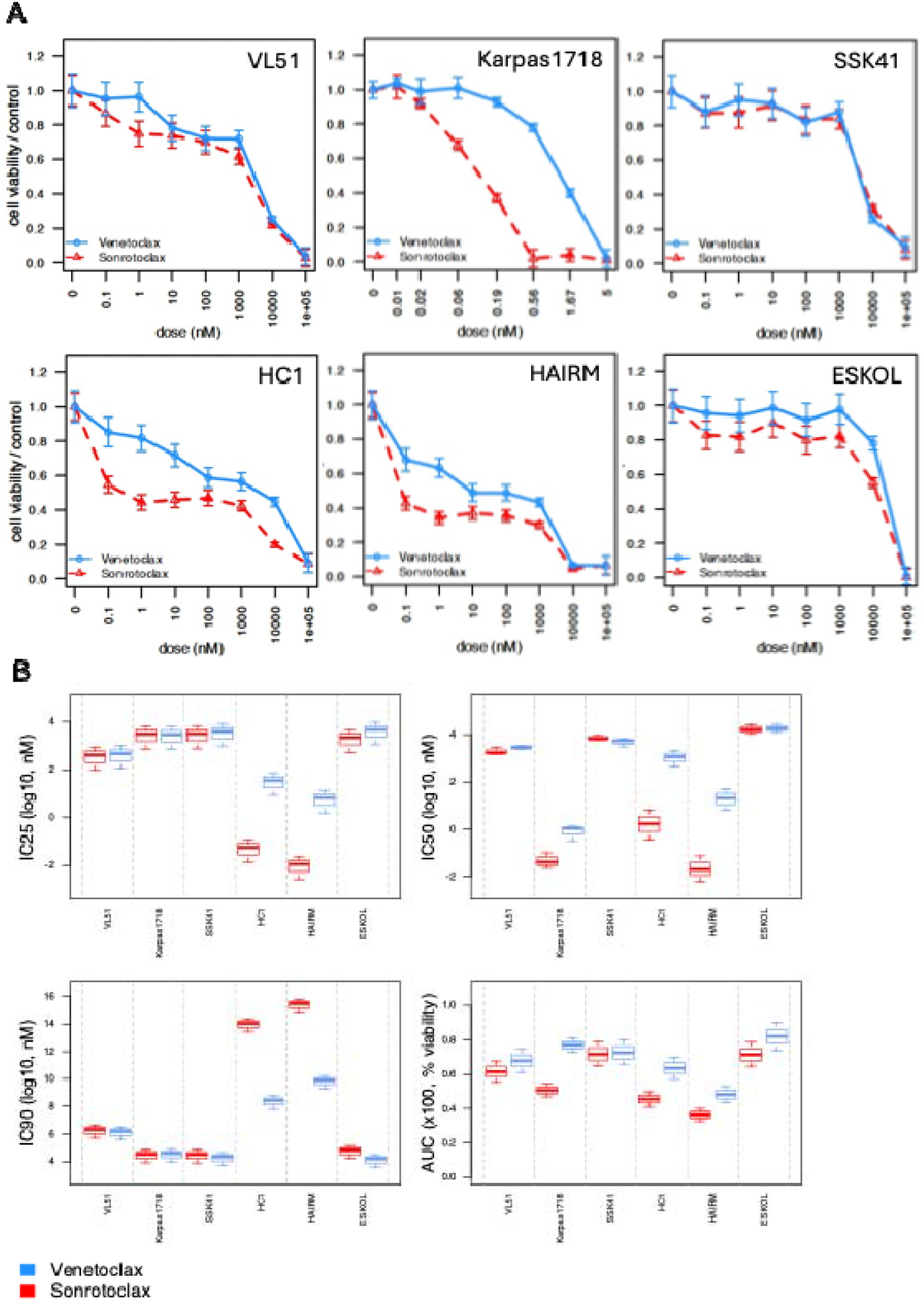
Effect of five days of exposure to the BCL2 inhibitors sonrotoclax and venetoclax in six MZL cell lines. (A) Drug response curves, (B) Boxplots with sensitivity parameters (AUC for Area under the curve). MTT after 5 days of exposure. Red, sonrotoclax. blue, venetoclax.

### The binding mode of BGB-16673 to BTK

We elucidated the binding mode of BGB-16673 to BTK via docking calculations, which is a widely employed technique to study ligand/protein binding interactions ^54-58^. We first investigated the binding of BGB-16673 to the kinase domain of human BTK (UniProt ID: Q06187, residues 395– 656), taking advantage of the reported crystallographic atomic structure of the protein in complex with the BTK inhibitor 4-∼{tert}-butyl-∼{N}-[2-methyl-3-[6-[4-(4-methylpiperazin-1-yl)carbonylphenyl]-7∼{H}-pyrrolo[2,3-d]pyrimidin-4-yl]phenyl]benzamide ^59^, which we refer to hereafter as L0Z (PDB ID: 6S90). Among the top-scoring docking poses, we identified a binding mode that occupies the BTK active site similarly to zanubrutinib (Figure 6A) and closely resembles the experimental binding conformation of L0Z, a known BTK inhibitor co-crystallized with the kinase domain (Figure 6B). The binding poses of BGB-16673 and L0Z showed a low root mean square deviation (r.m.s.d.) of 0.8□Å when calculated over their structurally conserved moiety. In both compounds, the tert-butyl group is oriented toward TYR551 and nestled within a hydrophobic pocket formed by residues such as PHE413, LEU542, and VAL546.

**Figure 6.**
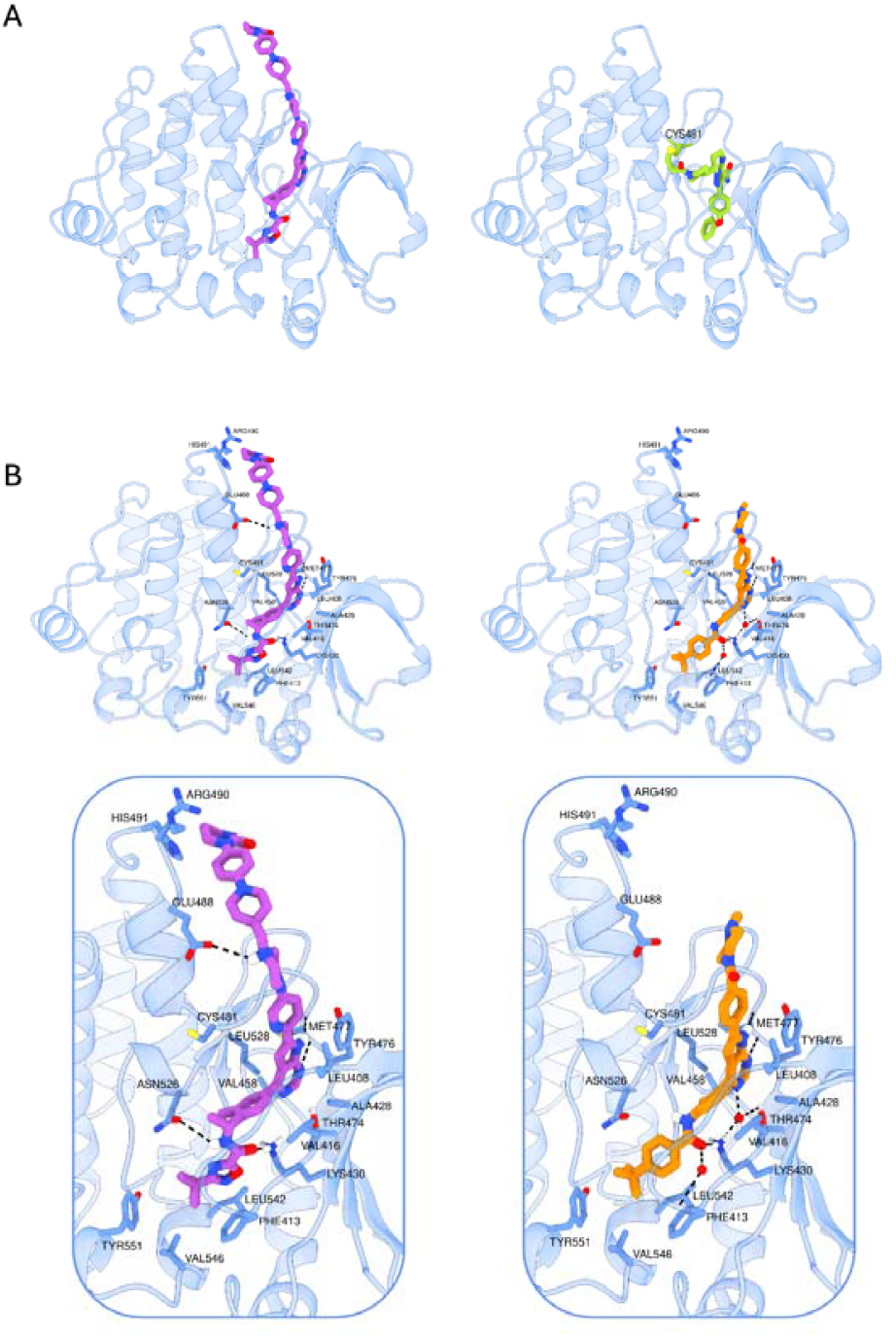
BGB-16673 binding mode to human BTK. A) Binding mode of BGB-16673 (purple sticks) to BTK predicted by docking calculations. The crystallographic binding mode of zanubrutinib (PDB ID: 6J6M) (green sticks) to BTK. The BTK protein is shown as light blue ribbons with the Cys481 residue covalently bound to zanubrutinib represented as green sticks. B) Three-dimensional representation of the binding conformation of BGB-16673 predicted by docking calculations and the crystallographic binding mode of the BTK-inhibitor L0Z (PDB ID: 6S90) within the BTK binding site. The protein is depicted as light blue ribbons, with side chains of interacting residues highlighted as blue sticks; BGB-16673 and L0Z are shown in purple and orange sticks, respectively. Protein–ligand interactions are illustrated as black dashed lines. For clarity, only hydrogen atoms directly involved in interactions are displayed.

The nearby carbonyl group of the amide in BGB-16673 forms a hydrogen bond with the side chain of LYS430 and may participate in a water-mediated interaction with the backbone of PHE413, as observed in the crystal structure of the L0Z–BTK complex. Additionally, BGB-16673 establishes a direct hydrogen bond between its amide group and the side chain of ASN526.

The pyrrolopyrimidine moiety in both ligands occupies a hydrophobic pocket formed by LEU408, VAL416, ALA428, VAL458, TYR476, and LEU528. Within this pocket, the pyrrole and pyrimidine rings form direct hydrogen bonds with the backbone of MET477. The pyrimidine ring may also engage in a water-mediated interaction with the side chains of LYS430 and THR474, consistent with interactions seen in the L0Z–BTK complex.

Unlike L0Z, BGB-16673 forms an additional strong salt bridge between the protonated amine of its piperazine moiety and the side chain of GLU488. Moreover, the terminal portion of BGB-16673 is positioned to potentially interact with the side chains of ARG490 and HIS491, providing further stabilization of the binding mode. A superposition of the binding conformations of BGB-16673 and L0Z is shown in Supplementary Figure 10.

## Discussion

Here, we showed that the BTK-degrader BGB-16673 has single-agent activity in individual MZL and MCL cell lines, it synergizes with other targeted agents in MZL cell lines, and its pattern of activity is like the BTK inhibitor zanubrutinib. We also report that the second-generation BCL2 inhibitor sonrotoclax showed higher activity than the first-generation BCL2 inhibitor venetoclax in MZL cell lines.

The BTK degrader BGB-16673 showed potent single-agent activity, particularly low IC50 values, in two MZL and two MCL cell lines. These findings are consistent with the essential role of BTK in B-cell malignancies ^1-4,60^. They also underscore the efficacy of targeted BTK degradation as a therapeutic strategy, which is now undergoing early clinical evaluation ^20,23,25-27,61^.

In the MZL models, we studied and in our experimental conditions, we did not observe differences between the degradation and the simple inhibition of BTK, indicating that the target selectivity is maintained in BGB-16673. Primary resistance to BGB-16673 is associated with preserved BTK phosphorylation and elevated p-BTK/BTK ratios, despite similar levels of total BTK degradation.

The effect on p-BTK inhibition was comparable, despite that the degradation of the whole protein was exclusively induced by BGB-16673 and not by zanubrutinib. We cannot exclude that the latter phenomenon might differentiate the activity of the degrader from the inhibitor in other experimental settings in the clinical context. Indeed, in line with this, RNA-Seq analysis also revealed a substantial overlap in the transcriptomic changes induced by BGB-16673 and zanubrutinib, but it also pinpointed possible differences between the two compounds. The significant downregulation of MYC targets and B-cell receptor signaling genes underscores the shared pathways targeted by BTK inhibition and degradation in these cellular models. BGB-16673 uniquely downregulated transcripts involved in oxidative phosphorylation, which could be a potential differentiator in its mechanism of action and therapeutic effect. Notably, oxidative phosphorylation is a metabolic process that we and others have associated with resistance to anti-cancer agents, including BTK inhibitors ^62-66^. This effect and the removal of the whole protein might lead to a reduced risk of resistance compared to BTK inhibitors when lymphoma cells are exposed to the compounds for a long time. However, this must be validated in future preclinical and clinical studies.

Molecular docking studies further showed that BGB-16673 occupies the BTK active site similarly to zanubrutinib, adopting a binding mode closely resembling the reported crystallographic binding mode of the non-covalent selective inhibitor L0Z ^59^.

Preclinical evidence indicates that BGB-16673 and other BTK-degraders in clinical development can overcome the resistance driven by BTK mutations ^21,22,24,67,68^. Interestingly, Montoya et al. demonstrated that the BTK and IKZF1/3 degrader NX-2127 can also counteract the oncogenic scaffold function of mutant BTK proteins, which activate additional kinases such as HCK and ILK, contributing to resistance against BTK inhibitors ^61^. We also tested BGB-16673 in the MZL model, which does not bear *BTK* mutations, with secondary resistance to BTK or PI3K inhibitors^15,16,34-36^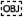. ^15,16,34,36^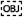. The BGB-16673 activity was reduced in this model, excluding the role of BTK as a scaffold to sustain the active signaling cascades that support the resistance in these models.

The synergistic potential of BGB-16673 was notably pronounced when combined with other agents, particularly BCL2 inhibitors (venetoclax and sonrotoclax), lenalidomide, rituximab, and bendamustine. The enhanced apoptotic response observed highlights a promising avenue for combination therapies with BGB-16673 in MZL. The observed benefit of combining BGB-16673 and lenalidomide is noteworthy since both compounds exploit cereblon to degrade their targets32,69.

The superior efficacy of sonrotoclax, a second-generation BCL2 inhibitor with reported more robust preclinical anti-tumor activity in a few models derived from MCL and diffuse large B cell lymphoma ^30^, over the first-generation BCL2 inhibitor venetoclax in certain cell lines is particularly noteworthy, suggesting it is a potent agent in MZL as a single agent or a partner in combination regimens. Indeed, despite the promising results achieved with a manageable safety profile and durable responses in a subset of patients treated with venetoclax ^6^, BCL2 inhibitors have not been exploited in MZL patients so far. Our data show higher activity than venetoclax in MZL models, and the early results from the ongoing phase 1 trials in patients with MZL ^33^ or Waldenström’s macroglobulinemia ^70^ strongly support the clinical development of this agent for MZL patients, as a single or in combination. The preliminary results reported for 14 R/R MZL patients enrolled in a phase 2 study combining ibrutinib and venetoclax support the dual BTK and BCL2 inhibition for this disease as a safe and active regimen with a complete remission rate of 43% according to PET/CT response at week 56 ^10,12^. Data obtained in patients with chronic lymphocytic leukemia indicate that also combining sonrotoclax and zanubrutinib also seems safe ^71^. No data are available with sonrotoclax combined with BTK degraders, but the ongoing trial NCT06634589 explores the combination of sonrotoclax and BGB-16673 in patients with R/R B-cell lymphomas.

Overall, our results affirm the potent and specific activity of BGB-16673 against BTK in MZL and MCL cell lines. Its ability to degrade BTK protein, combined with its synergistic effects with other targeted therapies, positions BGB-16673 as a promising candidate for further clinical development in MZL patients. Our study also identifies the second-generation BCL2 inhibitor sonrotoclax as another drug to be explored in the same patient populations, possibly combining the two molecules.

## Author contribution

AJA performed experiments, analyzed and interpreted data, performed data mining, prepared the figures, and co-wrote the manuscript.

LC performed data mining.

EC, CS: performed experiments.

LR, SR, VL: performed modeling computational studies. EZ, DR: provided advice.

AR performed transcriptome profiling.

FB co-designed research, interpreted data, and co-wrote the manuscript. All authors reviewed and accepted the final version of the manuscript.

## References

1. Burger JA, Wiestner A. Targeting B cell receptor signalling in cancer: preclinical and clinical advances. Nat Rev Cancer. 2018;18(3):148–167.

2. McDonald C, Xanthopoulos C, Kostareli E. The role of Bruton’s tyrosine kinase in the immune system and disease. Immunology. 2021;164(4):722–736.

3. Stephens DM, Byrd JC. Resistance to Bruton tyrosine kinase inhibitors: the Achilles heel of their success story in lymphoid malignancies. Blood. 2021;138(13):1099–1109.

4. Rossi D, Bertoni F, Zucca E. Marginal-Zone Lymphomas. N Engl J Med. 2022;386(6):568–581.

5. Tam C, Thompson PA. BTK inhibitors in CLL: second-generation drugs and beyond. Blood Adv. 2024;8(9):2300–2309.

6. Davids MS, Roberts AW, Kenkre VP, et al. Long-term Follow-up of Patients with Relapsed or Refractory Non-Hodgkin Lymphoma Treated with Venetoclax in a Phase I, First-in-Human Study. Clin Cancer Res. 2021;27(17):4690–4695.

7. Noy A, de Vos S, Thieblemont C, et al. Targeting Bruton tyrosine kinase with ibrutinib in relapsed/refractory marginal zone lymphoma. Blood. 2017;129(16):2224–2232.

8. Opat S, Tedeschi A, Hu B, et al. Safety and efficacy of zanubrutinib in relapsed/refractory marginal zone lymphoma: final analysis of the MAGNOLIA study. Blood Adv. 2023;7(22):6801–6811.

9. Deng L, Li Z, Zhang H, et al. Orelabrutinib for the treatment of relapsed or refractory marginal zone lymphoma: A phase 2, multicenter, open-label study. Am J Hematol. 2023;98(11):1742–1750.

10. Handunnetti SM, Khot A, Anderson MA, et al. Safety and Efficacy of Ibrutinib in Combination with Venetoclax in Patients with Marginal Zone Lymphoma: Preliminary Results from an Open Label, Phase II Study. Blood. 2019;134(Supplement_1):3999–3999.

11. Tarantelli C, Lange M, Gaudio E, et al. Copanlisib synergizes with conventional and targeted agents including venetoclax in B- and T-cell lymphoma models. Blood Adv. 2020;4(5):819–829.

12. Davis JE, L. LH, Ludford-Menting M, et al. Marginal Zone Lymphoma Is Associated with Dysregulated Peripheral Blood Immunity Which Is Not Changed By Ibrutinib-Venetoclax Treatment. Blood. 2024;144(Supplement 1):4384–4384.

13. de Vos S, Swinnen LJ, Wang D, et al. Venetoclax, bendamustine, and rituximab in patients with relapsed or refractory NHL: a phase Ib dose-finding study. Ann Oncol. 2018;29(9):1932–1938.

14. Nakhoda S, Vistarop A, Wang YL. Resistance to Bruton tyrosine kinase inhibition in chronic lymphocytic leukaemia and non-Hodgkin lymphoma. Br J Haematol. 2023;200(2):137–149.

15. Arribas AJ, Napoli S, Cascione L, et al. ERBB4-Mediated Signaling Is a Mediator of Resistance to PI3K and BTK Inhibitors in B-cell Lymphoid Neoplasms. Mol Cancer Ther. 2024;23(3):368–380.

16. Arribas AJ, Napoli S, Cascione L, et al. Resistance to PI3Kdelta inhibitors in marginal zone lymphoma can be reverted by targeting the IL-6/PDGFRA axis. Haematologica. 2022;107(11):2685–2697.

17. Stephens DM, Byrd JC. Next-Generation Bruton Tyrosine Kinase Inhibitors. J Clin Oncol. 2020;38(25):2937–2940.

18. Lue JK, O’Connor OA, Bertoni F. Targeting pathogenic mechanisms in marginal zone lymphoma: from concepts and beyond. Ann Lymphoma. 2020;4:7.

19. Feng X, Wang Y, Long T, et al. P1239: Bruton tyrosine kinase (BTK) protein degrader BGB-16673 is less apt to cause, and able to overcome variable BTK resistance mutations compared to other BTK inhibitors (BTKi). Hemasphere. 2023;7(Suppl).

20. Danilov A, Tees MT, Patel K, et al. A First-in-Human Phase 1 Trial of NX-2127, a First-in-Class Bruton’s Tyrosine Kinase (BTK) Dual-Targeted Protein Degrader with Immunomodulatory Activity, in Patients with Relapsed/Refractory B Cell Malignancies. Blood. 2023;142(Supplement 1):4463–4463.

21. Robbins DW, Noviski MA, Tan YS, et al. Discovery and Preclinical Pharmacology of NX-2127, an Orally Bioavailable Degrader of Bruton’s Tyrosine Kinase with Immunomodulatory Activity for the Treatment of Patients with B Cell Malignancies. J Med Chem. 2024;67(4):2321–2336.

22. Noviski MA, Brathaban N, Mukerji R, et al. Abstract 2850: NX-5948 promotes selective, sub-nanomolar degradation of inhibitor-resistant BTK mutants. Cancer Res. 2023;83(7_Supplement):2850–2850.

23. Searle E, Forconi F, Linton K, et al. Initial Findings from a First-in-Human Phase 1a/b Trial of NX-5948, a Selective Bruton’s Tyrosine Kinase (BTK) Degrader, in Patients with Relapsed/Refractory B Cell Malignancies. Blood. 2023;142(Supplement 1):4473–4473.

24. Wang H, Hou X, Zhang W, et al. P1219: BGB-16673, a BTK degrader, overcomes on-target resistance from BTK inhibitors and presents sustainable long-term tumor regression in lymphoma xenograft models. Hemasphere. 2023;7(Suppl).

25. Tam CS, Frustaci AM, Bijou F, et al. Preliminary Efficacy and Safety of the Bruton Tyrosine Kinase Degrader BGB-16673 in Patients with Relapsed or Refractory (R/R) Indolent NHL: Results from the Phase 1 CaDAnCe-101 Study. Blood. 2024;144(Supplement 1):1649–1649.

26. Seymour JF, Tam CS, Cheah CY, et al. Preliminary Efficacy and Safety of the Bruton Tyrosine Kinase Degrader BGB-16673 in Patients with Relapsed or Refractory Waldenström Macroglobulinemia: Results from the Phase 1 CaDAnCe-101 Study. Blood. 2024;144(Supplement 1):860–860.

27. Shah NN, Omer Z, Collins GP, et al. Efficacy and Safety of the Bruton’s Tyrosine Kinase (BTK) Degrader NX-5948 in Patients with Relapsed/Refractory (R/R) Chronic Lymphocytic Leukemia (CLL): Updated Results from an Ongoing Phase 1a/b Study. Blood. 2024;144(Supplement 1):884–884.

28. Seymour JF, Cheah CY, Parrondo R, et al. First Results from a Phase 1, First-in-Human Study of the Bruton’s Tyrosine Kinase (BTK) Degrader Bgb-16673 in Patients (Pts) with Relapsed or Refractory (R/R) B-Cell Malignancies (BGB-16673-101). Blood. 2023;142:4401.

29. Zinzani PL, Frustaci AM, Narkhede M, et al. 436 | UPDATED EFFICACY/SAFETY OF BRUTON TYROSINE KINASE DEGRADER BGB-16673 IN PATIENTS WITH RELAPSED/REFRACTORY INDOLENT NHL: ONGOING PHASE 1 CaDAnCe-101 RESULTS. Hematol Oncol. 2025;43(S3):e436.70094.

30. Liu J, Li S, Wang Q, et al. Sonrotoclax overcomes BCL2 G101V mutation-induced venetoclax resistance in preclinical models of hematologic malignancy. Blood. 2024;143(18):1825–1836.

31. Guo Y, Xue H, Hu N, et al. Discovery of the Clinical Candidate Sonrotoclax (BGB-11417), a Highly Potent and Selective Inhibitor for Both WT and G101V Mutant Bcl-2. J Med Chem. 2024;67(10):7836–7858.

32. Wu Y, Meibohm B, Zhang T, et al. Translational modelling to predict human pharmacokinetics and pharmacodynamics of a Bruton’s tyrosine kinase-targeted protein degrader BGB-16673. Br J Pharmacol. 2024;181(24):4973–4987.

33. Tedeschi A, Cheah CY, Opat SS, et al. Monotherapy with Second-Generation BCL2 Inhibitor Sonrotoclax (BGB-11417) Is Well Tolerated with High Response Rates in Patients with Relapsed/Refractory Marginal Zone Lymphoma: Data from an Ongoing Phase 1 Study. Blood. 2023;142(Supplement 1):3032–3032.

34. Arribas A, Napoli S, Cascione L, et al. Secondary resistance to the PI3K inhibitor copanlisib in marginal zone lymphoma. Eur J Cancer. 2020;138:S40–S40.

35. Arribas AJ, Cascione L, Cannas E, et al. Abstract 534: Novel marginal zone lymphoma models of resistance to the dual inhibition of PI3K and BCL2. Cancer Res. 2024;84(6_Supplement):534–534.

36. Arribas AJ, Guidetti F, Cannas E, et al. IL-16 production is a mechanism of resistance to BTK inhibitors and R-CHOP in lymphomas. bioRxiv. 2025.

37. Johnson Z, Tarantelli C, Civanelli E, et al. IOA-244 is a Non-ATP-competitive, Highly Selective, Tolerable PI3K Delta Inhibitor That Targets Solid Tumors and Breaks Immune Tolerance. Cancer Res Commun. 2023;3(4):576–591.

38. Boi M, Gaudio E, Bonetti P, et al. The BET Bromodomain Inhibitor OTX015 Affects Pathogenetic Pathways in Preclinical B-cell Tumor Models and Synergizes with Targeted Drugs. Clin Cancer Res. 2015;21(7):1628–1638.

39. Smirnov P, Safikhani Z, El-Hachem N, et al. PharmacoGx: an R package for analysis of large pharmacogenomic datasets. Bioinformatics. 2016;32(8):1244–1246.

40. Berenbaum MC. What is synergy? Pharmacol Rev. 1989;41(2):93–141.

41. Zheng S, Wang W, Aldahdooh J, et al. SynergyFinder Plus: Toward Better Interpretation and Annotation of Drug Combination Screening Datasets. Genomics Proteomics Bioinformatics. 2022;20(3):587–596.

42. Andrews S. FastQC A Quality Control tool for High Throughput Sequence Data; 2010.

43. Bolger AM, Lohse M, Usadel B. Trimmomatic: a flexible trimmer for Illumina sequence data. Bioinformatics. 2014;30(15):2114–2120.

44. Dobin A, Davis CA, Schlesinger F, et al. STAR: ultrafast universal RNA-seq aligner. Bioinformatics. 2013;29(1):15–21.

45. Anders S, Pyl PT, Huber W. HTSeq--a Python framework to work with high-throughput sequencing data. Bioinformatics. 2015;31(2):166–169.

46. Robinson MD, McCarthy DJ, Smyth GK. edgeR: a Bioconductor package for differential expression analysis of digital gene expression data. Bioinformatics. 2010;26(1):139–140.

47. Law CW, Chen Y, Shi W, Smyth GK. voom: Precision weights unlock linear model analysis tools for RNA-seq read counts. Genome Biol. 2014;15(2):R29.

48. Subramanian A, Tamayo P, Mootha VK, et al. Gene set enrichment analysis: a knowledge-based approach for interpreting genome-wide expression profiles. Proc Natl Acad Sci U S A. 2005;102(43):15545–15550.

49. Shaffer AL, Wright G, Yang L, et al. A library of gene expression signatures to illuminate normal and pathological lymphoid biology. Immunol Rev. 2006;210:67–85.

50. O’Boyle NM, Banck M, James CA, Morley C, Vandermeersch T, Hutchison GR. Open Babel: An open chemical toolbox. J Cheminform. 2011;3:33.

51. Pan X, Wang H, Li C, Zhang JZH, Ji C. MolGpka: A Web Server for Small Molecule pK(a) Prediction Using a Graph-Convolutional Neural Network. J Chem Inf Model. 2021;61(7):3159–3165.

52. Pettersen EF, Goddard TD, Huang CC, et al. UCSF ChimeraX: Structure visualization for researchers, educators, and developers. Protein Sci. 2021;30(1):70–82.

53. Morris GM, Huey R, Lindstrom W, et al. AutoDock4 and AutoDockTools4: Automated docking with selective receptor flexibility. J Comput Chem. 2009;30(16):2785–2791.

54. Limongelli V. Ligand binding free energy and kinetics calculation in 2020. Wiley Interdisciplinary Reviews-Computational Molecular Science. 2020;10(4):e1455.

55. Di Leva FS, Di Marino D, Limongelli V. Structural Insight into the Binding Mode of FXR and GPBAR1 Modulators. Handb Exp Pharmacol. 2019;256:111–136.

56. Heckmann D, Laufer B, Marinelli L, et al. Breaking the dogma of the metal-coordinating carboxylate group in integrin ligands: introducing hydroxamic acids to the MIDAS to tune potency and selectivity. Angew Chem Int Ed Engl. 2009;48(24):4436–4440.

57. Di Leva FS, Festa C, Renga B, et al. Structure-based drug design targeting the cell membrane receptor GPBAR1: exploiting the bile acid scaffold towards selective agonism. Sci Rep. 2015;5:16605.

58. Carino A, Biagioli M, Marchiano S, et al. Disruption of TFGbeta-SMAD3 pathway by the nuclear receptor SHP mediates the antifibrotic activities of BAR704, a novel highly selective FXR ligand. Pharmacol Res. 2018;131:17–31.

59. Pulz R, Angst D, Dawson J, et al. Design of Potent and Selective Covalent Inhibitors of Bruton’s Tyrosine Kinase Targeting an Inactive Conformation. ACS Med Chem Lett. 2019;10(10):1467–1472.

60. Bonfiglio F, Bruscaggin A, Guidetti F, et al. Genetic and phenotypic attributes of splenic marginal zone lymphoma. Blood. 2022;139(5):732–747.

61. Montoya S, Bourcier J, Noviski M, et al. Kinase-impaired BTK mutations are susceptible to clinical-stage BTK and IKZF1/3 degrader NX-2127. Science. 2024;383(6682):eadi5798.

62. Zhao Z, Mei Y, Wang Z, He W. The Effect of Oxidative Phosphorylation on Cancer Drug Resistance. Cancers (Basel). 2022;15(1).

63. Tarantelli C, Gaudio E, Arribas AJ, et al. PQR309 Is a Novel Dual PI3K/mTOR Inhibitor with Preclinical Antitumor Activity in Lymphomas as a Single Agent and in Combination Therapy. Clin Cancer Res. 2018;24(1):120–129.

64. Liu Y, Kimpara S, Hoang NM, et al. EGR1-mediated metabolic reprogramming to oxidative phosphorylation contributes to ibrutinib resistance in B-cell lymphoma. Blood. 2023;142(22):1879–1894.

65. Zammarchi F, Havenith KE, Sachini N, et al. ADCT-602, a Novel PBD Dimer-containing Antibody-Drug Conjugate for Treating CD22-positive Hematologic Malignancies. Mol Cancer Ther. 2024;23(4):520–531.

66. Mensah AA, Spriano F, Sartori G, et al. Study of the antilymphoma activity of pracinostat reveals different sensitivities of DLBCL cells to HDAC inhibitors. Blood Adv. 2021;5(10):2467–2480.

67. Pan C, Riehm J, Gururaja TL, et al. Abstract 605: ABBV-101, a highly potent and selective clinical stage Bruton tyrosine kinase degrader, overcomes BTK mutation-induced resistance to BTK inhibitors. Cancer Res. 2024;84(6_Supplement):605–605.

68. Li J, Xu W, Yan P, Cao Y, Hu M, Daley W. Abstract CT128: Phase 1 study of HSK29116, a Bruton tyrosine kinase (BTK) proteolysis-targeting chimera (PROTAC) agent, in patients with relapsed or refractory B-cell malignancies. Cancer Res. 2023;83(8_Supplement):CT128–CT128.

69. Lopez-Girona A, Mendy D, Ito T, et al. Cereblon is a direct protein target for immunomodulatory and antiproliferative activities of lenalidomide and pomalidomide. Leukemia. 2012;26(11):2326–2335.

70. Cheah C, Tam C, Sanz RG, et al. P1110: Safety and efficacy results of a phase 1 study of the novel BCL2 inhibitor sonrotoclax (BGB-11417) for relapsed/refractory Waldenström’s macroglobulinemia. Hemasphere. 2024;8(S1):2017–2018.

71. Soumerai JD, Cheah CY, Anderson MA, et al. Sonrotoclax and Zanubrutinib as Frontline Treatment for CLL Demonstrates High MRD Clearance Rates with Good Tolerability: Data from an Ongoing Phase 1/1b Study BGB-11417-101. Blood. 2024;144(Supplement 1):1012–1012.

