## Supplementary figures and tables for "Dual targeting of BTK and BCL2 enhances apoptosis in marginal zone lymphoma models: preclinical activity of BGB-16673 and sonrotoclax"

**Supplementary Figure 1.** Effect of exposure to the BTK inhibitors ibrutinib and zanubrutinib, or the BTK-degrader BGB-16673 in MCL and MZL cell lines. (A) Effect of exposure to the BTK inhibitors Ibrutinib or Zanubrutinib in two MCL cell lines, REC1 (black) and MINO (red). (B) Effect of exposure to the BTK-degrader BGB-16673 (blue) or to the BTK inhibitor Zanubrutinib (orange) in two MZL cell lines, Karpas1718 (left) and SSK41 (right)
MTT assay after five days of exposure.

A

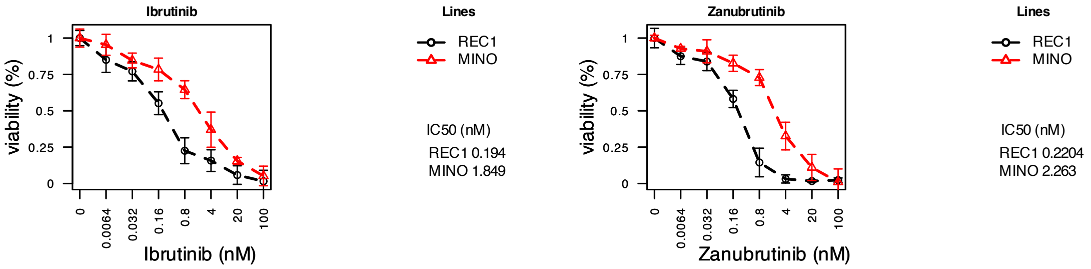


B

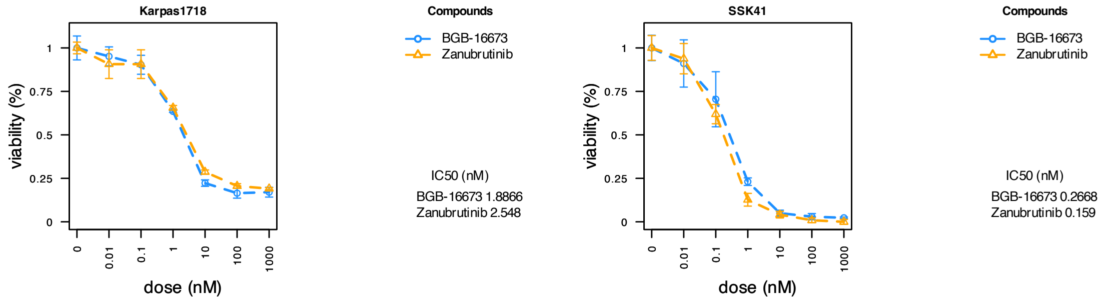

**Supplementary Figure 2.** Effect of the BTK-degrader BGB-16673 and the BTK inhibitor zanubrutinib on the protein levels of pBTK and BTK in the two sensitive MZL cell lines Karpas1718 (24hr, A) and SSK41 (24hr, B; 3hr and 6hr C); and in the four primary resistant MZL cell lines: VL51 (3hr, D), ESKOL (3hr, E), HAIRM, (3hr, F) and HC1 (3hr, G). Immunoblotting is representative of two independent experiments. Each barplot shows the mean of the protein quantification in the two experiments, error bars correspond to the standard deviation of the mean, and numbers are the p-value from a t-test comparing each treatment to control (DMSO, grey bar). H) Protein levels of BTK, pBTK and pBTK/BTK (top and left middle panels, blue) upon BGB-16673, and IC50 values (right panel, red), were compared between sensitive (Karpas1718 and SSK41) and primary resistant (VL51, ESKOL, HAIRM and HC1) cell lines by t-test. The bottom panel shows a heatmap on the protein levels of BTK, pBTK and pBTK/BTK across cell lines tested.

A) Karpas1718

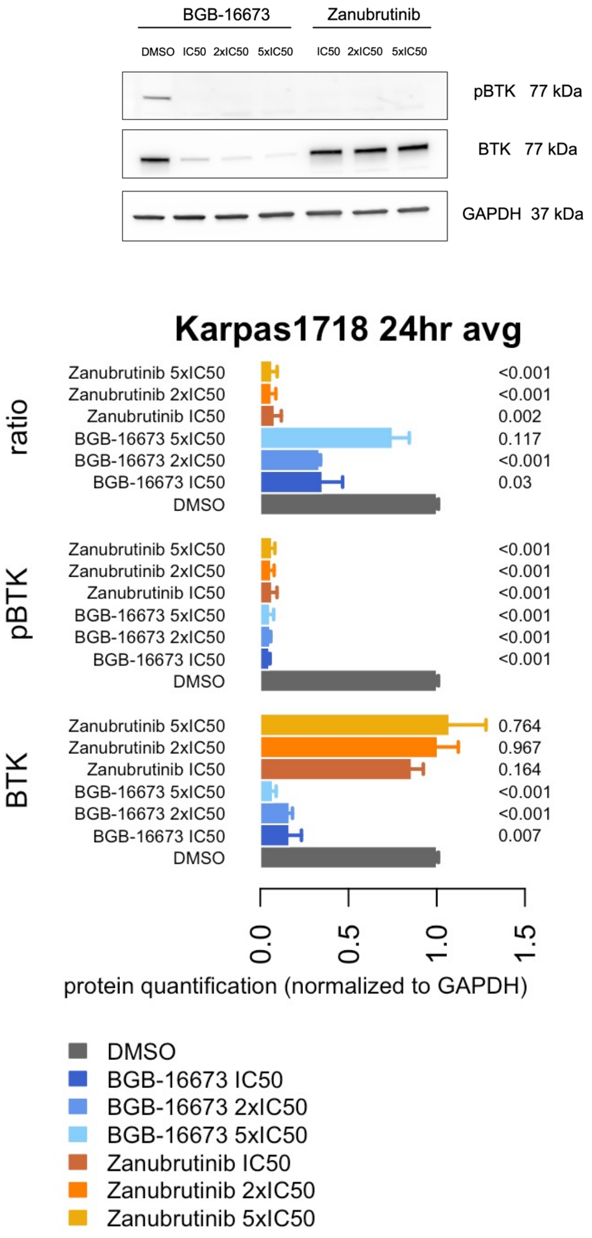

B)

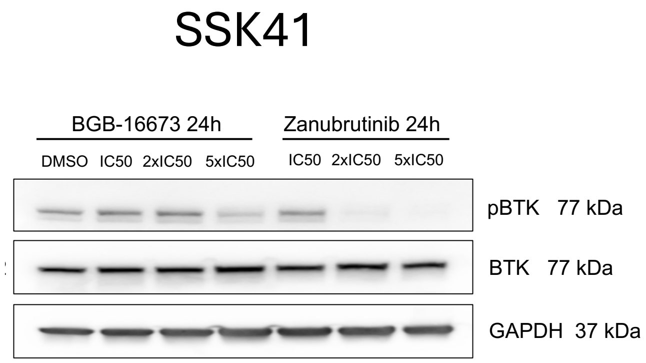
SSK41
BGB-16673 24h
Zanubrutinib 24h
DMSO
IC50
2xIC50
5xIC50
IC50
2xIC50
5xIC50
pBTK 77 kDa
BTK 77 kDa
GAPDH 37 kDa

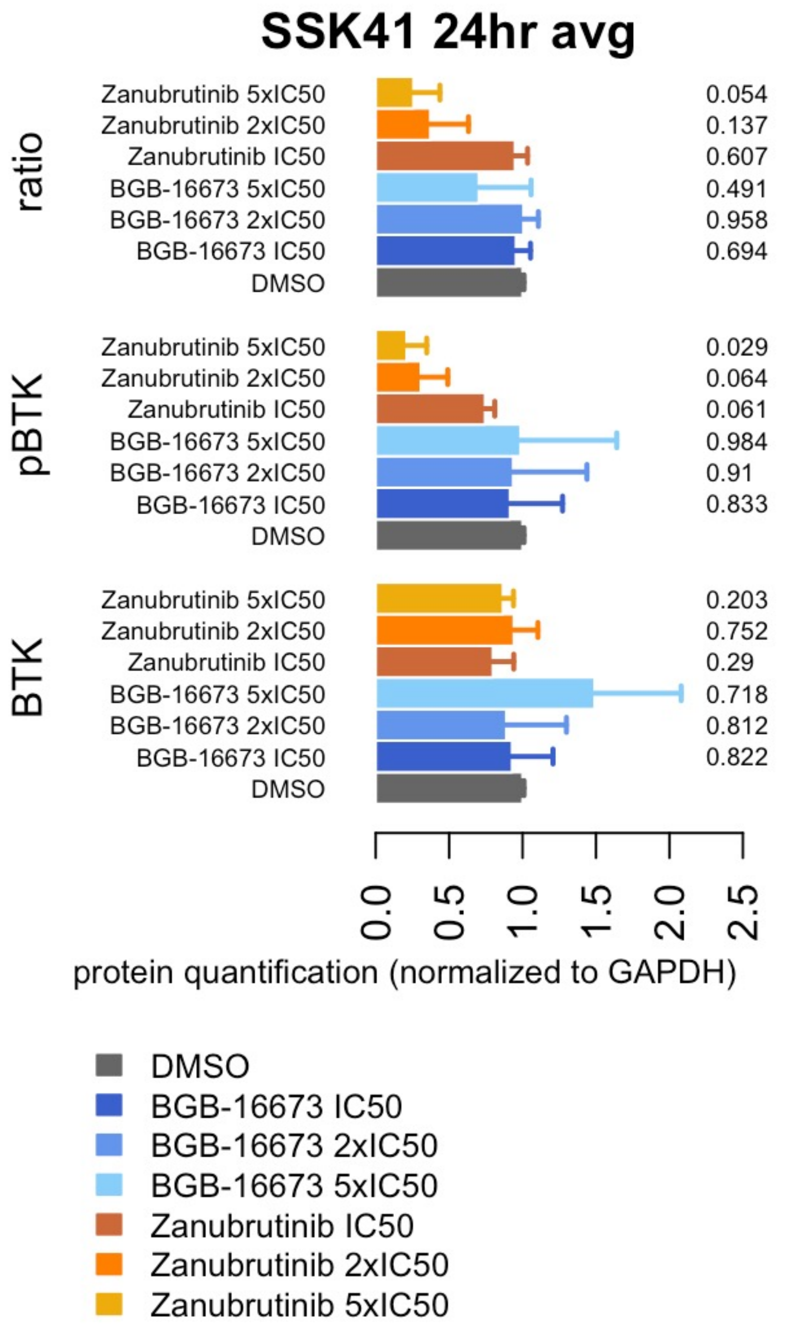
SSK41 24hr avg
ratio
Zanubrutinib 5xIC50 0.054
Zanubrutinib 2xIC50 0.137
Zanubrutinib IC50 0.607
BGB-16673 5xIC50 0.491
BGB-16673 2xIC50 0.958
BGB-16673 IC50 0.694
DMSO
pBTK
Zanubrutinib 5xIC50 0.029
Zanubrutinib 2xIC50 0.064
Zanubrutinib IC50 0.061
BGB-16673 5xIC50 0.984
BGB-16673 2xIC50 0.91
BGB-16673 IC50 0.833
DMSO
BTK
Zanubrutinib 5xIC50 0.203
Zanubrutinib 2xIC50 0.752
Zanubrutinib IC50 0.29
BGB-16673 5xIC50 0.718
BGB-16673 2xIC50 0.812
BGB-16673 IC50 0.822
DMSO
0.0
0.5
1.0
1.5
2.0
2.5
protein quantification (normalized to GAPDH)
DMSO
BGB-16673 IC50
BGB-16673 2xIC50
BGB-16673 5xIC50
Zanubrutinib IC50
Zanubrutinib 2xIC50
Zanubrutinib 5xIC50

C)

SSK41

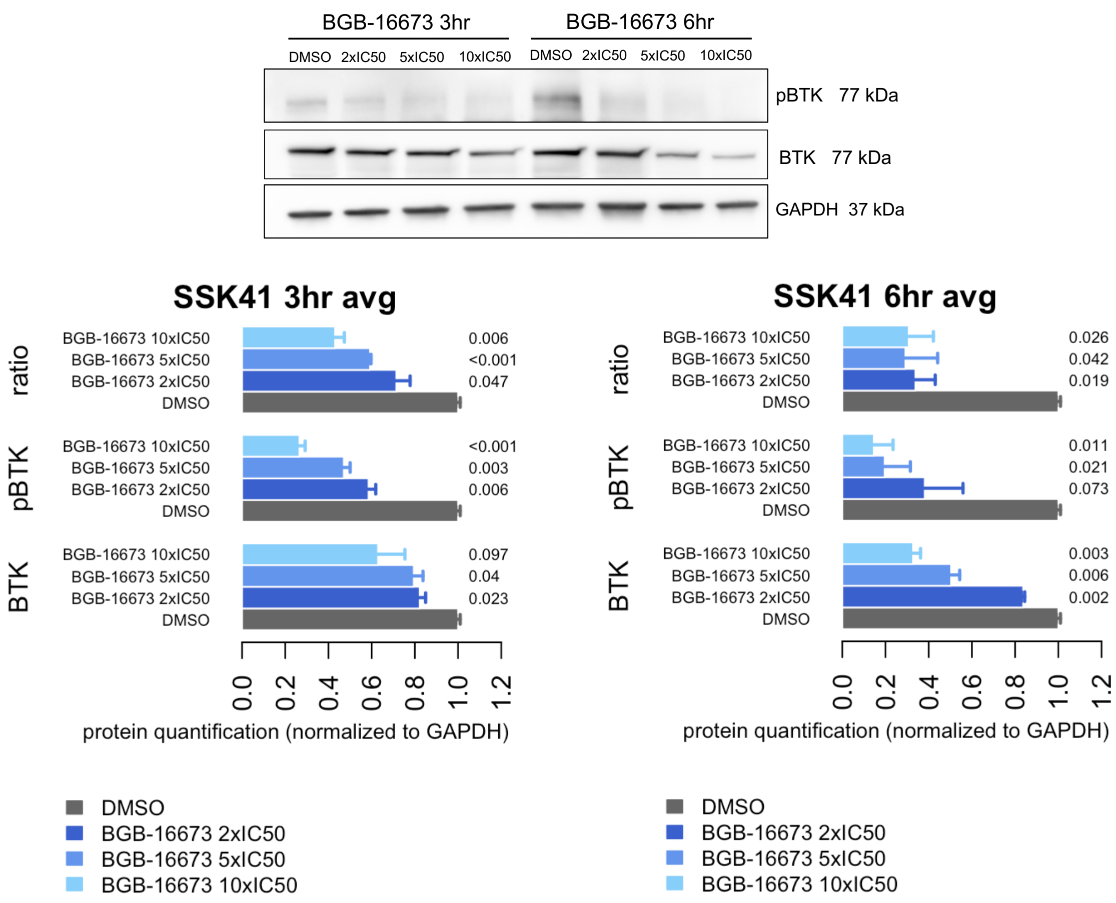

BGB-16673 3hr
BGB-16673 6hr
DMSO 2xIC50 5xIC50 10xIC50 DMSO 2xIC50 5xIC50 10xIC50
pBTK 77 kDa
BTK 77 kDa
GAPDH 37 kDa
SSK41 3hr avg
SSK41 6hr avg
ratio
pBTK
BTK
BGB-16673 10xIC50
BGB-16673 5xIC50
BGB-16673 2xIC50
DMSO
0.006
<0.001
0.047
<0.001
0.003
0.006
0.097
0.04
0.023
0.026
0.042
0.019
0.011
0.021
0.073
0.003
0.006
0.002
0.0 0.2 0.4 0.6 0.8 1.0 1.2
protein quantification (normalized to GAPDH)
DMSO
BGB-16673 2xIC50
BGB-16673 5xIC50
BGB-16673 10xIC50

D) VL51

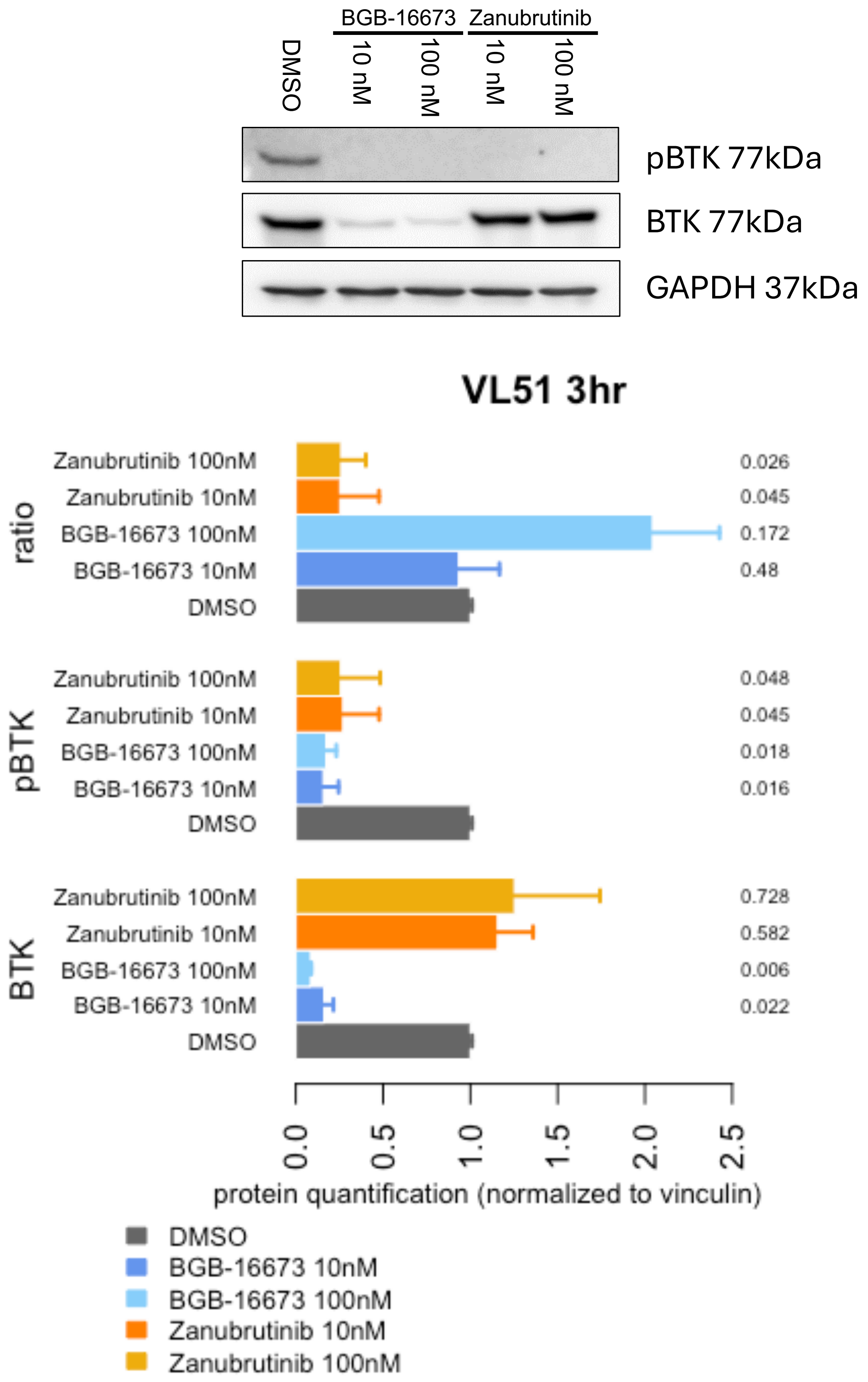

BGB-16673
Zanubrutinib
DMSO
10 nM
100 nM
10 nM
100 nM
pBTK 77kDa
BTK 77kDa
GAPDH 37kDa
VL51 3hr
ratio
pBTK
BTK
Zanubrutinib 100nM
Zanubrutinib 10nM
BGB-16673 100nM
BGB-16673 10nM
DMSO
0.026
0.045
0.172
0.48
0.048
0.045
0.018
0.016
0.728
0.582
0.006
0.022
0.0
0.5
1.0
1.5
2.0
2.5
protein quantification (normalized to vinculin)
DMSO
BGB-16673 10nM
BGB-16673 100nM
Zanubrutinib 10nM
Zanubrutinib 100nM

E) ESKOL

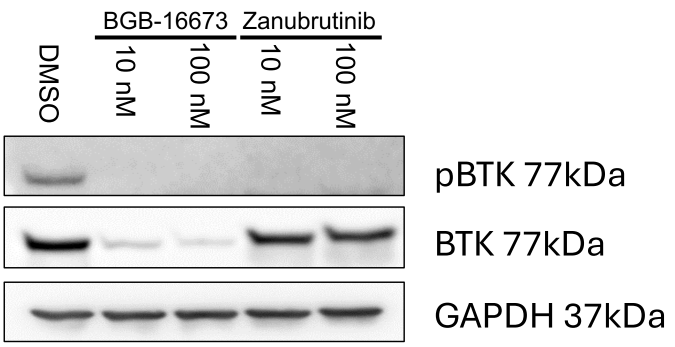

BGB-16673
Zanubrutinib
DMSO
10 nM
100 nM
10 nM
100 nM
pBTK 77kDa
BTK 77kDa
GAPDH 37kDa


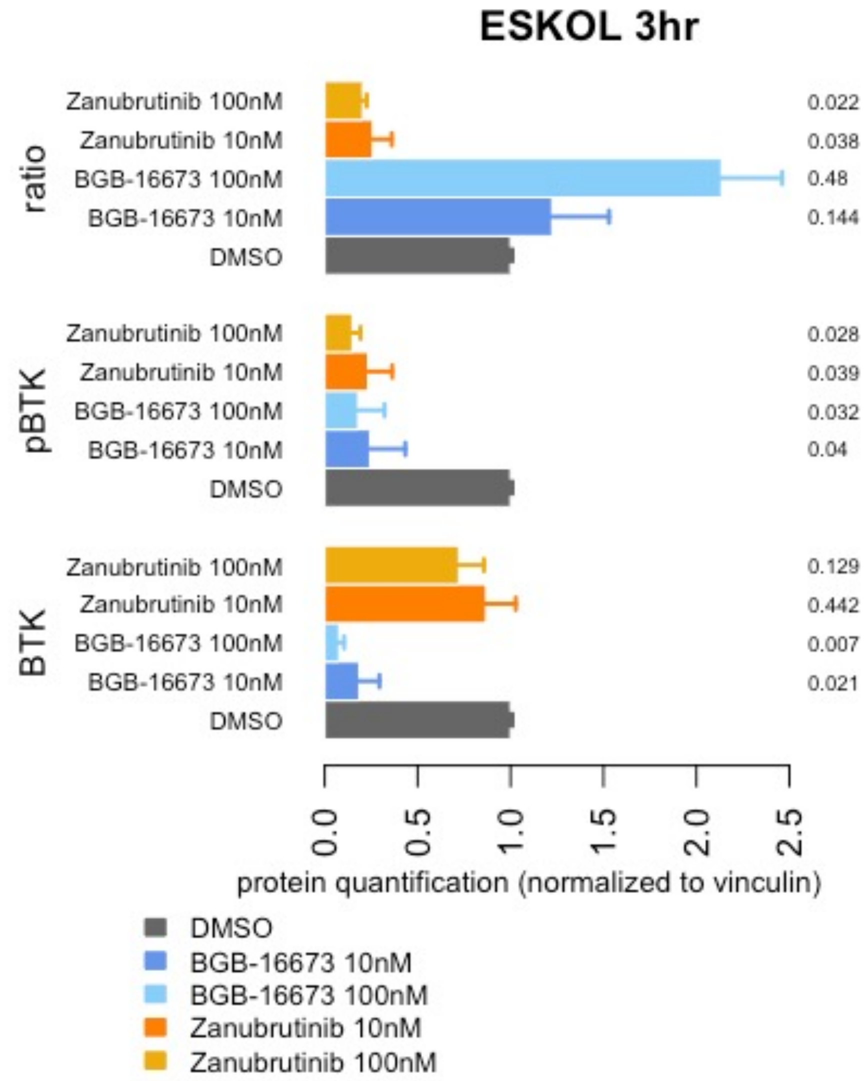

ESKOL 3hr
ratio
pBTK
BTK
Zanubrutinib 100nM
Zanubrutinib 10nM
BGB-16673 100nM
BGB-16673 10nM
DMSO
0.022
0.038
0.48
0.144
0.028
0.039
0.032
0.04
0.129
0.442
0.007
0.021
0.0
0.5
1.0
1.5
2.0
2.5
protein quantification (normalized to vinculin)
DMSO
BGB-16673 10nM
BGB-16673 100nM
Zanubrutinib 10nM
Zanubrutinib 100nM

F)

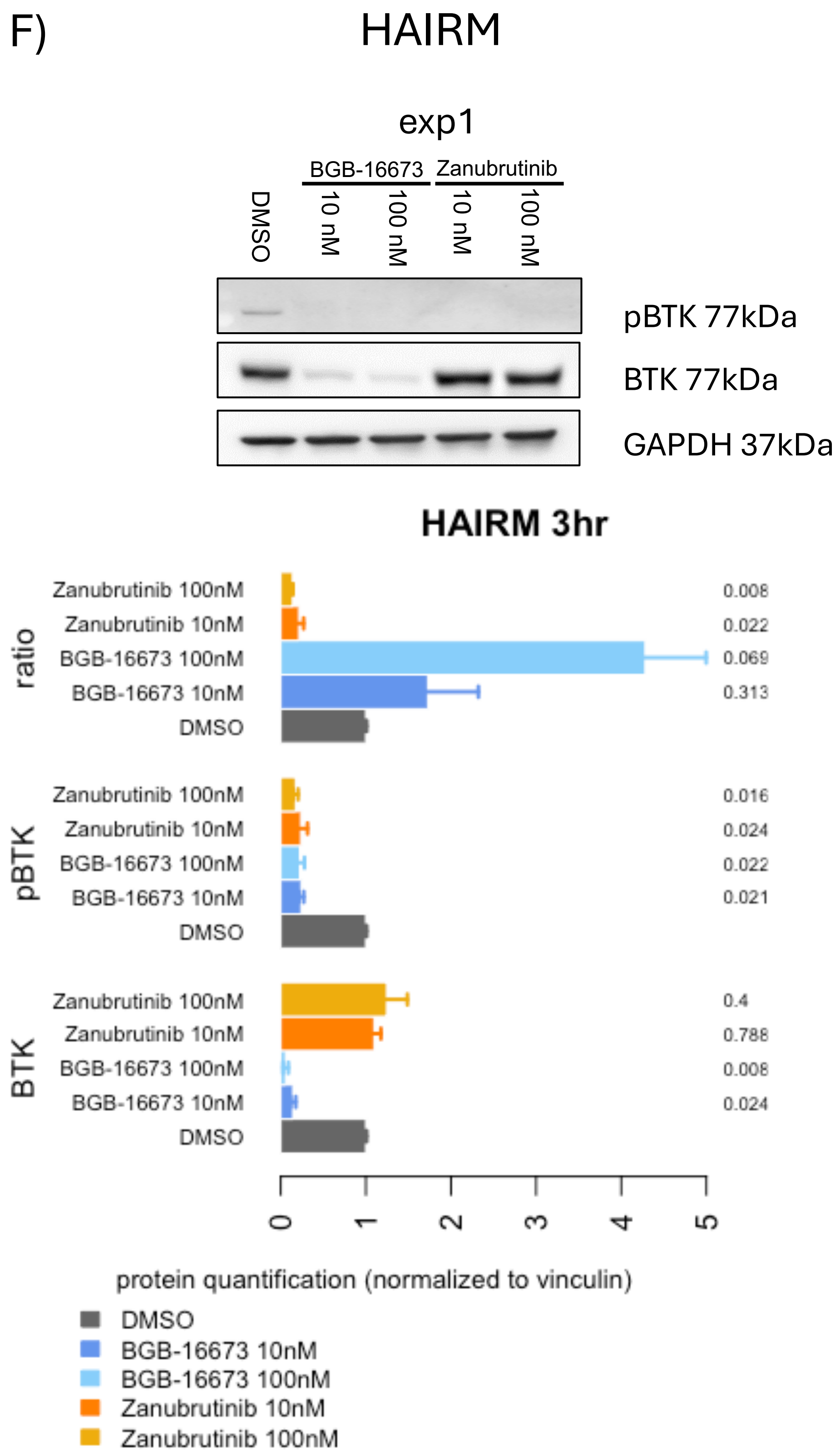

HAIRM
exp1
BGB-16673
Zanubrutinib
DMSO
10 nM
100 nM
10 nM
100 nM
pBTK 77kDa
BTK 77kDa
GAPDH 37kDa
HAIRM 3hr
ratio
pBTK
BTK
Zanubrutinib 100nM
Zanubrutinib 10nM
BGB-16673 100nM
BGB-16673 10nM
DMSO
0.008
0.022
0.069
0.313
0.016
0.024
0.022
0.021
0.4
0.788
0.008
0.024
0
1
2
3
4
5
protein quantification (normalized to vinculin)
DMSO
BGB-16673 10nM
BGB-16673 100nM
Zanubrutinib 10nM
Zanubrutinib 100nM

G) HC1

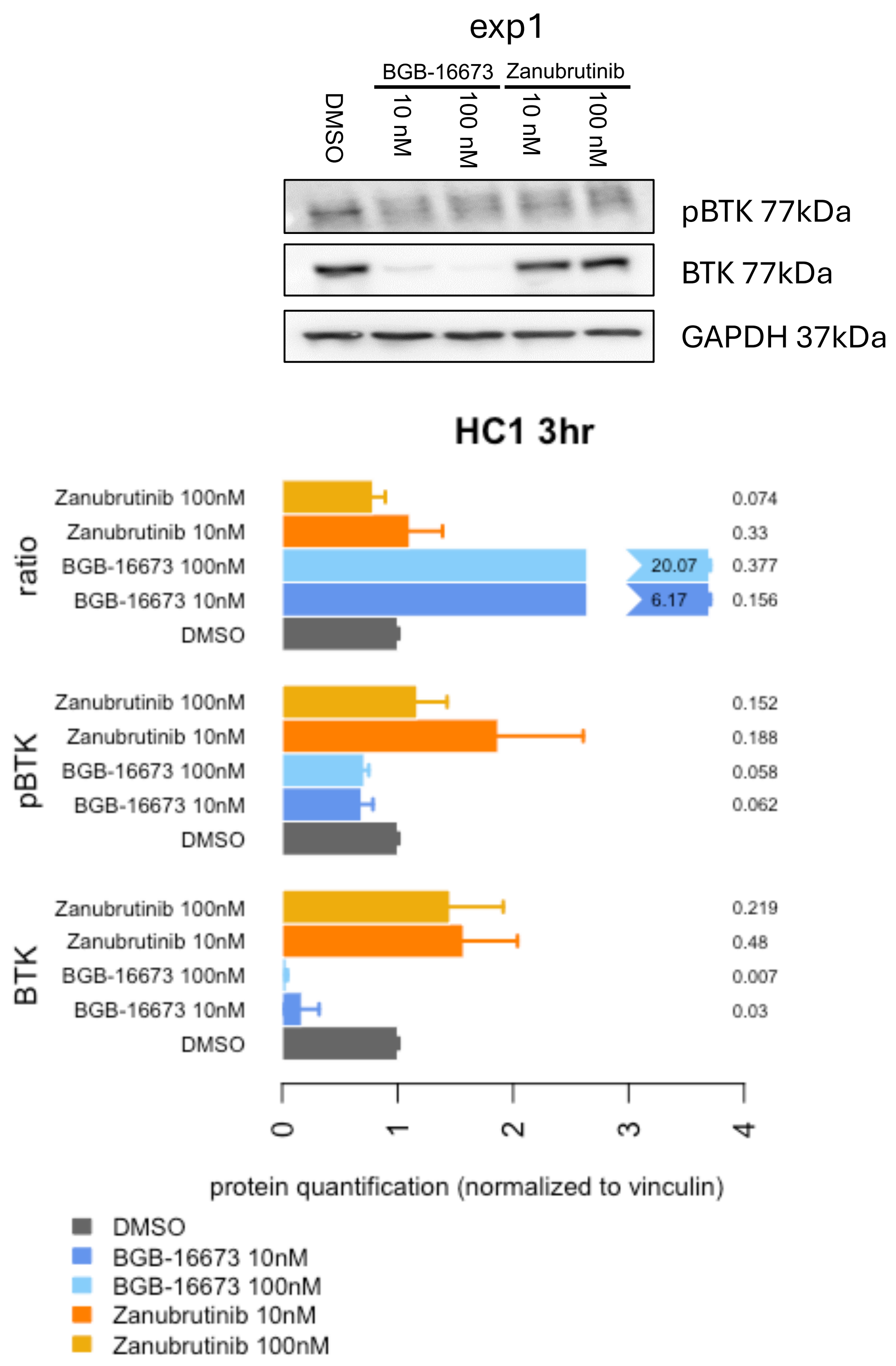
exp1
BGB-16673
Zanubrutinib
DMSO
10 nM
100 nM
10 nM
100 nM
pBTK 77kDa
BTK 77kDa
GAPDH 37kDa
HC1 3hr
ratio
pBTK
BTK
Zanubrutinib 100nM
Zanubrutinib 10nM
BGB-16673 100nM
BGB-16673 10nM
DMSO
20.07
6.17
0.074
0.33
0.377
0.156
0.152
0.188
0.058
0.062
0.219
0.48
0.007
0.03
0
1
2
3
4
protein quantification (normalized to vinculin)
DMSO
BGB-16673 10nM
BGB-16673 100nM
Zanubrutinib 10nM
Zanubrutinib 100nM

H)

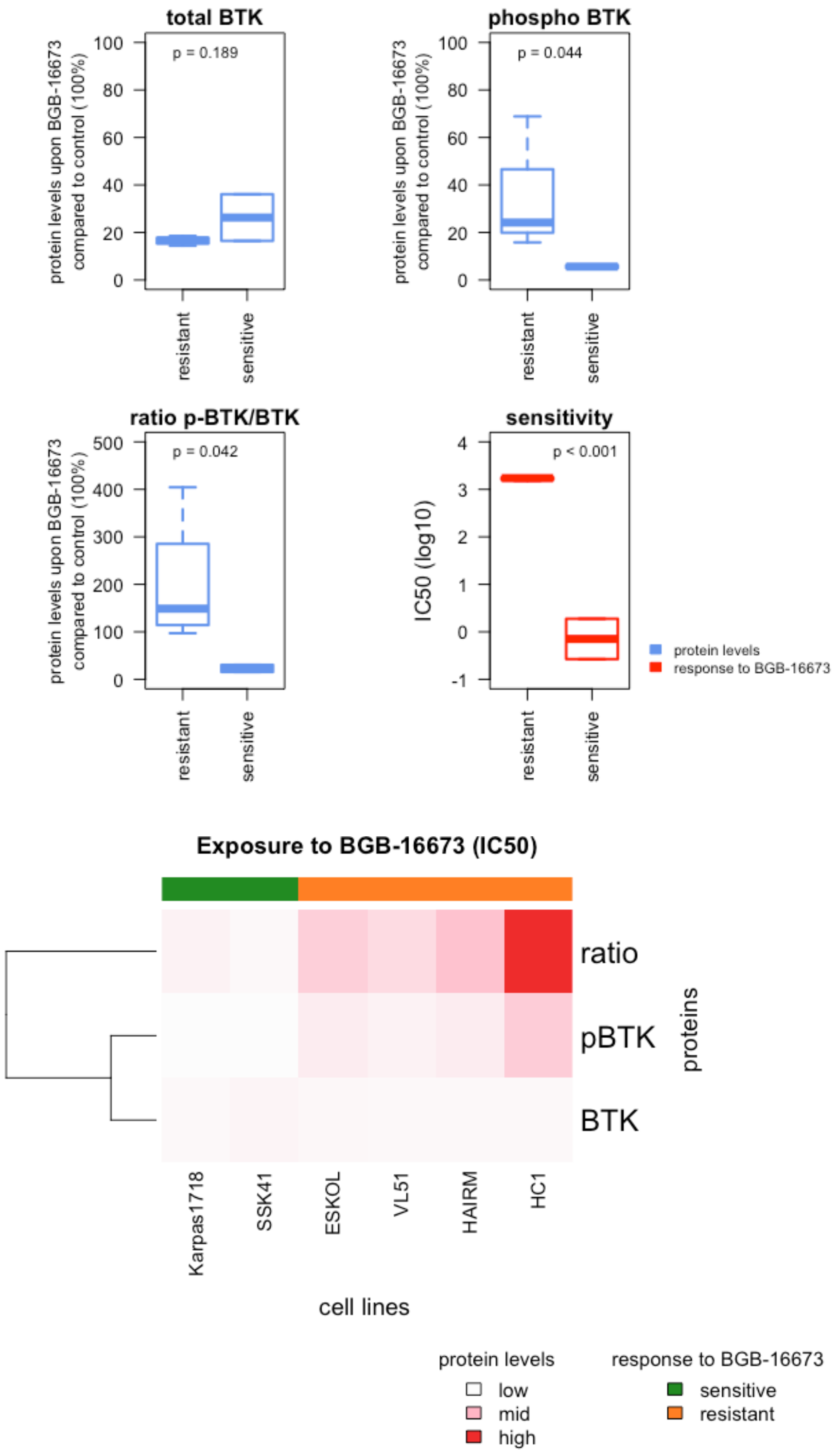

total BTK
p = 0.189
phospho BTK
p = 0.044
protein levels upon BGB-16673 compared to control (100%)
resistant
sensitive
ratio p-BTK/BTK
p = 0.042
sensitivity
p < 0.001
IC50 (log10)
protein levels
response to BGB-16673
Exposure to BGB-16673 (IC50)
ratio
pBTK
BTK
proteins
Karpas1718
SSK41
ESKOL
VL51
HAIRM
HC1
cell lines
protein levels
low
mid
high
response to BGB-16673
sensitive
resistant

**Supplementary Figure 3.** Effect of five days of exposure to the BTK-degrader BGB-16673 and of BGB-21704, a similar drug to BGB-16673 with no BTK-related activity, in the two MZL cell lines Karpas1718 (left panel) and SSK41 (right panel). MTT after 5 days of exposure. Blue, BGB-16673. grey, BGB-21704.

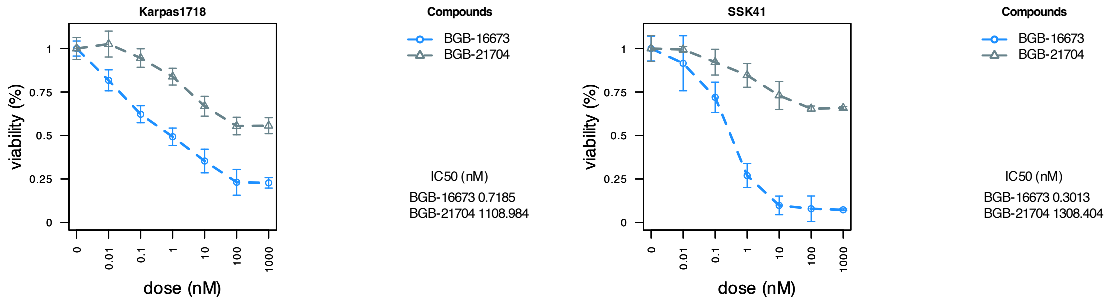

**Supplementary Figure 4.** Effect of five days of exposure to the BTK-degrader BGB-16673 in parental and resistant derivatives to PI3K, BTK, and BCL2 inhibitors, derived from Karpas1718 (top left), VL51 (top right) and SSK41 (bottom left). MTT after 5 days of exposure.

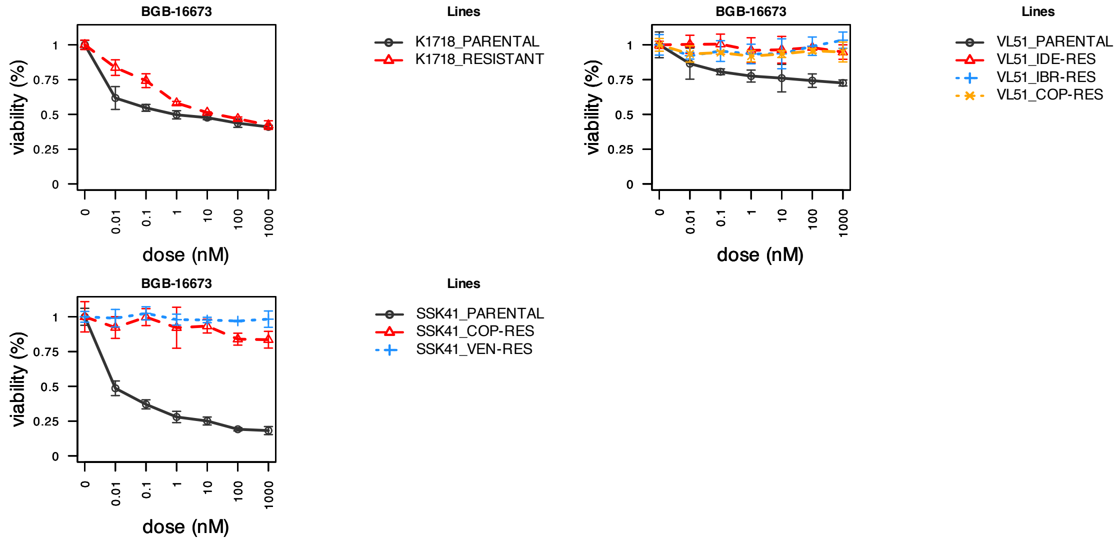

**Supplementary Figure 5.** Functional enrichment analyses from gene expression profiles of BGB-16673 or Zanubrutinib when compared to DMSO. Gene-concept network from Gene ontology enrichment: molecular function (A), cellular component (B) and biological process (C); from KEGG pathways (D for BGB-16673, E for zanubrutinib). Ridgeline plots of the distribution of the core-enriched genes of the top 10 significantly deregulated genesets from MSigDB Hallmarks and Reactome pathways of BGB-16673 (F and) or Zanubrutinib (G) expression profiles.

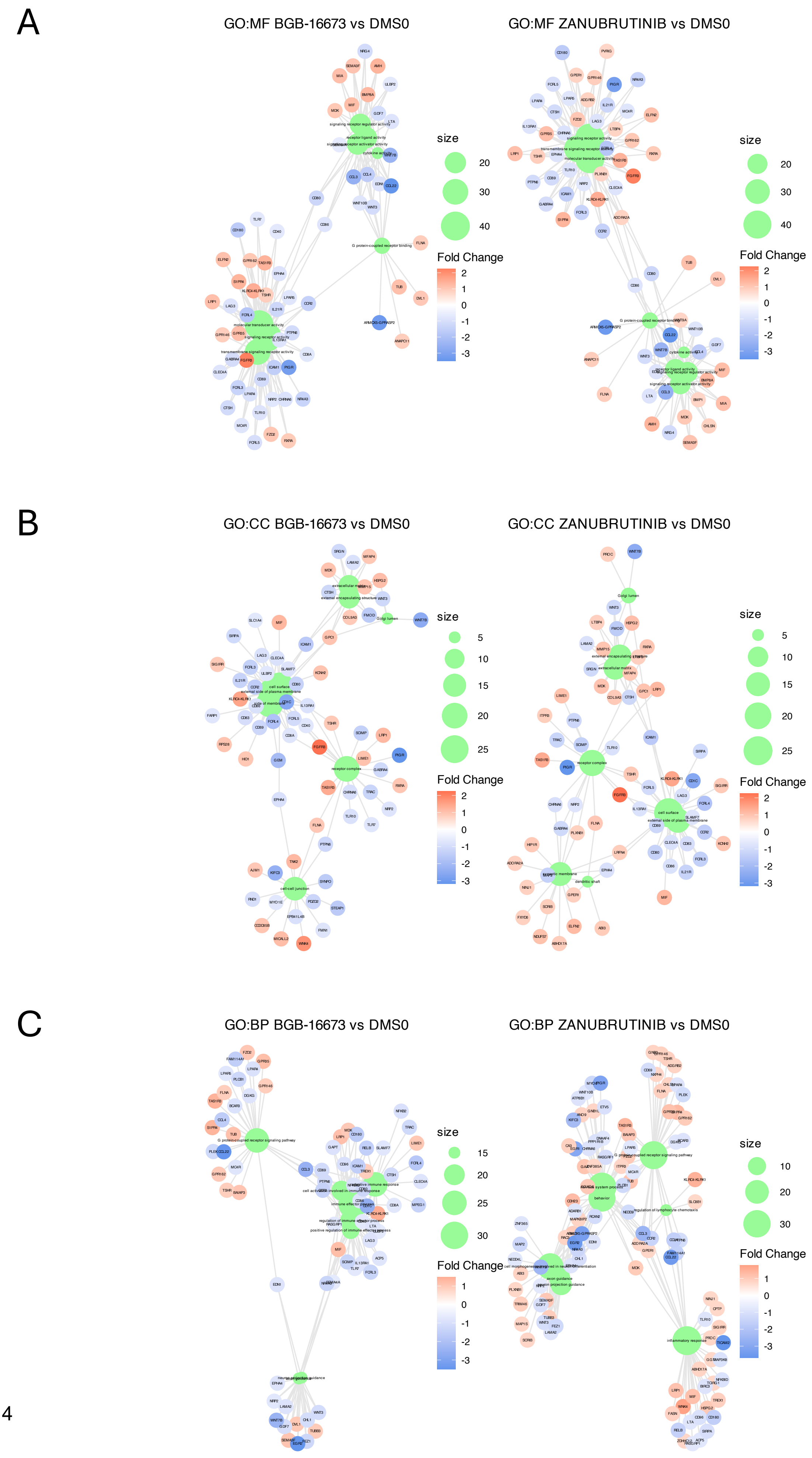
A
GO:MF BGB-16673 vs DMS0
GO:MF ZANUBRUTINIB vs DMS0
size
20
30
40
Fold Change
2
1
0
-1
-2
-3
B
GO:CC BGB-16673 vs DMS0
GO:CC ZANUBRUTINIB vs DMS0
size
5
10
15
20
25
Fold Change
2
1
0
-1
-2
-3
C
GO:BP BGB-16673 vs DMS0
GO:BP ZANUBRUTINIB vs DMS0
size
15
20
25
30
Fold Change
1
0
-1
-2
-3
size
10
20
30
Fold Change
1
0
-1
-2
-3

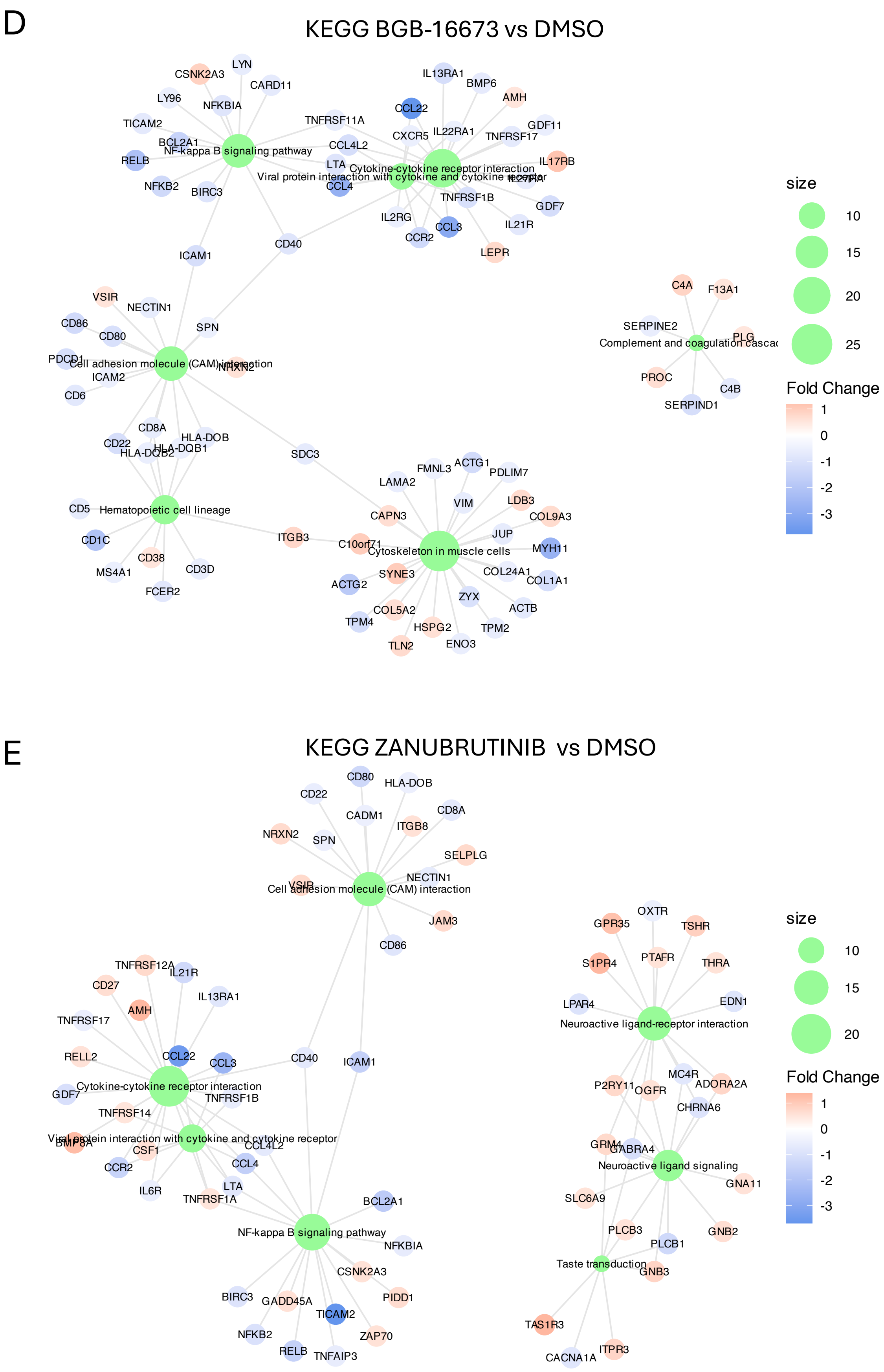
D
KEGG BGB-16673 vs DMSO
NF-kappa B signaling pathway
Cytokine-cytokine receptor interaction
Viral protein interaction with cytokine and cytokine receptor
Cell adhesion molecule (CAM) interaction
Hematopoietic cell lineage
Cytoskeleton in muscle cells
Complement and coagulation cascad
CSNK2A3
LYN
CARD11
LY96
NFKBIA
TICAM2
BCL2A1
RELB
NFKB2
BIRC3
TNFRSF11A
CCL4L2
LTA
CCL4
IL13RA1
BMP6
AMH
CCL22
CXCR5
IL22RA1
GDF11
TNFRSF17
IL17RB
IL27RA
TNFRSF1B
GDF7
IL2RG
CCL3
CCR2
IL21R
LEPR
CD40
ICAM1
VSIR
NECTIN1
CD86
CD80
SPN
PDCD1
NRXN2
ICAM2
CD6
CD8A
HLA-DOB
CD22
HLA-DQB2
HLA-DQB1
SDC3
CD5
CD1C
CD38
MS4A1
CD3D
FCER2
ITGB3
FMNL3
ACTG1
PDLIM7
LAMA2
VIM
LDB3
CAPN3
COL9A3
JUP
C10orf71
MYH11
SYNE3
COL24A1
ACTG2
COL1A1
ZYX
ACTB
COL5A2
TPM4
HSPG2
TPM2
TLN2
ENO3
C4A
F13A1
SERPINE2
PLG
PROC
C4B
SERPIND1
size
10
15
20
25
Fold Change
1
0
-1
-2
-3
E
KEGG ZANUBRUTINIB vs DMSO
Cell adhesion molecule (CAM) interaction
Cytokine-cytokine receptor interaction
Viral protein interaction with cytokine and cytokine receptor
NF-kappa B signaling pathway
Neuroactive ligand-receptor interaction
Neuroactive ligand signaling
Taste transduction
CD80
HLA-DOB
CD22
CADM1
CD8A
ITGB8
NRXN2
SPN
SELPLG
NECTIN1
VSIR
JAM3
CD86
TNFRSF12A
IL21R
CD27
IL13RA1
AMH
TNFRSF17
RELL2
CCL22
CCL3
CD40
ICAM1
GDF7
TNFRSF1B
TNFRSF14
BMP8A
CSF1
CCL4L2
CCR2
CCL4
IL6R
LTA
TNFRSF1A
BCL2A1
NFKBIA
CSNK2A3
BIRC3
GADD45A
PIDD1
TICAM2
NFKB2
ZAP70
RELB
TNFAIP3
OXTR
GPR35
TSHR
S1PR4
PTAFR
THRA
LPAR4
EDN1
MC4R
P2RY11
OGFR
ADORA2A
CHRNA6
GRM4
GABRA4
GNA11
SLC6A9
PLCB3
PLCB1
GNB2
GNB3
TAS1R3
ITPR3
CACNA1A
size
10
15
20
Fold Change
1
0
-1
-2
-3

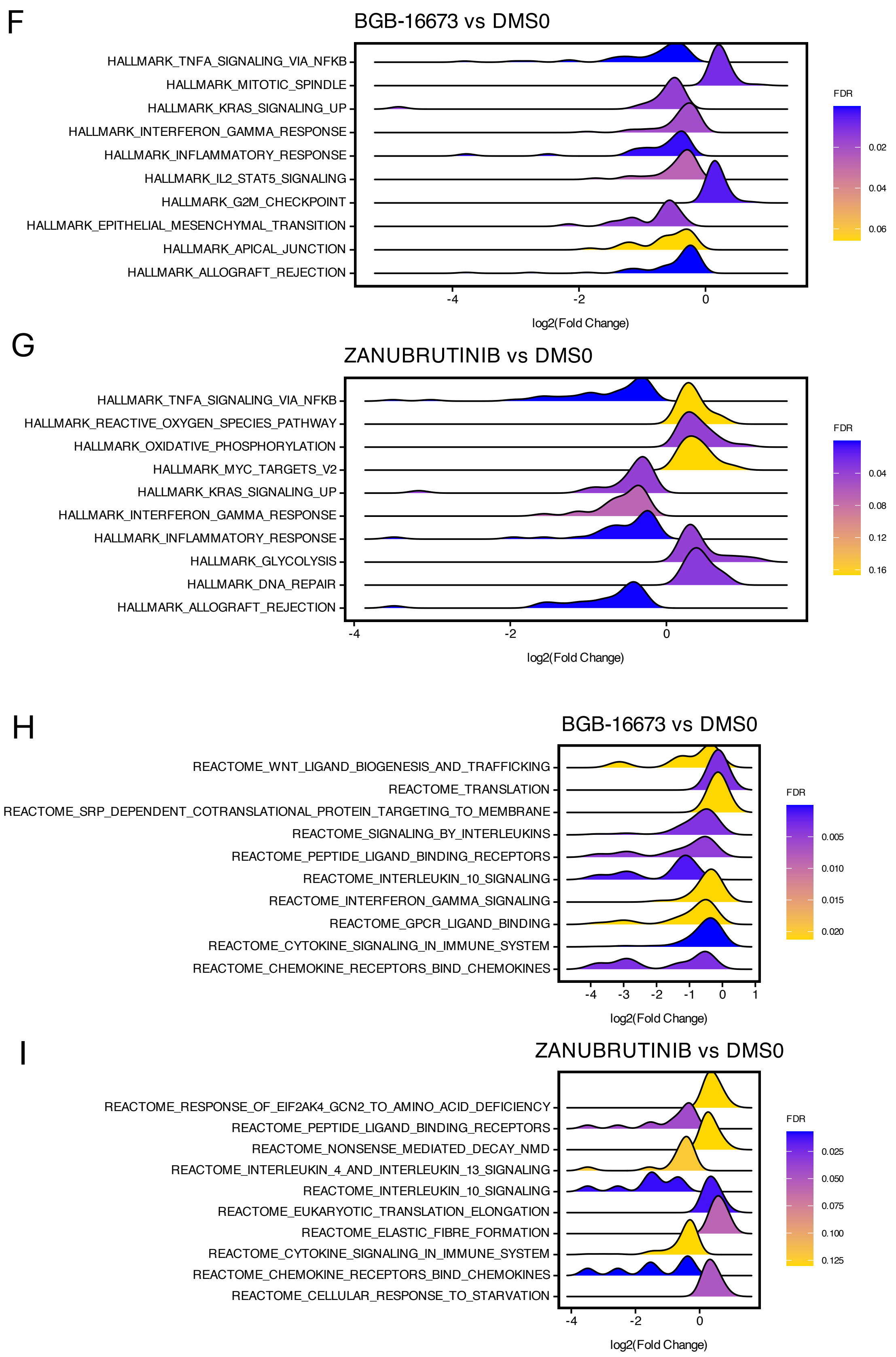
F
BGB-16673 vs DMS0
HALLMARK_TNFA_SIGNALING_VIA_NFKB
HALLMARK_MITOTIC_SPINDLE
HALLMARK_KRAS_SIGNALING_UP
HALLMARK_INTERFERON_GAMMA_RESPONSE
HALLMARK_INFLAMMATORY_RESPONSE
HALLMARK_IL2_STAT5_SIGNALING
HALLMARK_G2M_CHECKPOINT
HALLMARK_EPITHELIAL_MESENCHYMAL_TRANSITION
HALLMARK_APICAL_JUNCTION
HALLMARK_ALLOGRAFT_REJECTION
-4
-2
0
log2(Fold Change)
FDR
0.02
0.04
0.06
G
ZANUBRUTINIB vs DMS0
HALLMARK_TNFA_SIGNALING_VIA_NFKB
HALLMARK_REACTIVE_OXYGEN_SPECIES_PATHWAY
HALLMARK_OXIDATIVE_PHOSPHORYLATION
HALLMARK_MYC_TARGETS_V2
HALLMARK_KRAS_SIGNALING_UP
HALLMARK_INTERFERON_GAMMA_RESPONSE
HALLMARK_INFLAMMATORY_RESPONSE
HALLMARK_GLYCOLYSIS
HALLMARK_DNA_REPAIR
HALLMARK_ALLOGRAFT_REJECTION
-4
-2
0
log2(Fold Change)
FDR
0.04
0.08
0.12
0.16
H
BGB-16673 vs DMS0
REACTOME_WNT_LIGAND_BIOGENESIS_AND_TRAFFICKING
REACTOME_TRANSLATION
REACTOME_SRP_DEPENDENT_COTRANSLATIONAL_PROTEIN_TARGETING_TO_MEMBRANE
REACTOME_SIGNALING_BY_INTERLEUKINS
REACTOME_PEPTIDE_LIGAND_BINDING_RECEPTORS
REACTOME_INTERLEUKIN_10_SIGNALING
REACTOME_INTERFERON_GAMMA_SIGNALING
REACTOME_GPCR_LIGAND_BINDING
REACTOME_CYTOKINE_SIGNALING_IN_IMMUNE_SYSTEM
REACTOME_CHEMOKINE_RECEPTORS_BIND_CHEMOKINES
-4 -3 -2 -1 0 1
log2(Fold Change)
FDR
0.005
0.010
0.015
0.020
I
ZANUBRUTINIB vs DMS0
REACTOME_RESPONSE_OF_EIF2AK4_GCN2_TO_AMINO_ACID_DEFICIENCY
REACTOME_PEPTIDE_LIGAND_BINDING_RECEPTORS
REACTOME_NONSENSE_MEDIATED_DECAY_NMD
REACTOME_INTERLEUKIN_4_AND_INTERLEUKIN_13_SIGNALING
REACTOME_INTERLEUKIN_10_SIGNALING
REACTOME_EUKARYOTIC_TRANSLATION_ELONGATION
REACTOME_ELASTIC_FIBRE_FORMATION
REACTOME_CYTOKINE_SIGNALING_IN_IMMUNE_SYSTEM
REACTOME_CHEMOKINE_RECEPTORS_BIND_CHEMOKINES
REACTOME_CELLULAR_RESPONSE_TO_STARVATION
-4
-2
0
log2(Fold Change)
FDR
0.025
0.050
0.075
0.100
0.125

**Supplementary Figure 6.** Effect of the combination of BGB-16673 with a series of agents used in clinical settings or clinical development. Karpas1718, HC1, SSK41, and VL51 MZL cell lines were exposed to increasing concentrations of the BTK degrader and a series of clinically relevant agents for five days, followed by an MTT assay. The BTK degrader was combined with lenalidomide (A), selinexor (B), venetoclax (C), bendamustine (D), tazemetostat (E), and rituximab (F). The benefit of the combination was assessed according to the Highest Single Agent (HSA) algorithm in the Synergy Finder online tool (https://synergyfinder.aittokallio.group/). Red for synergistic, green for antagonistic.

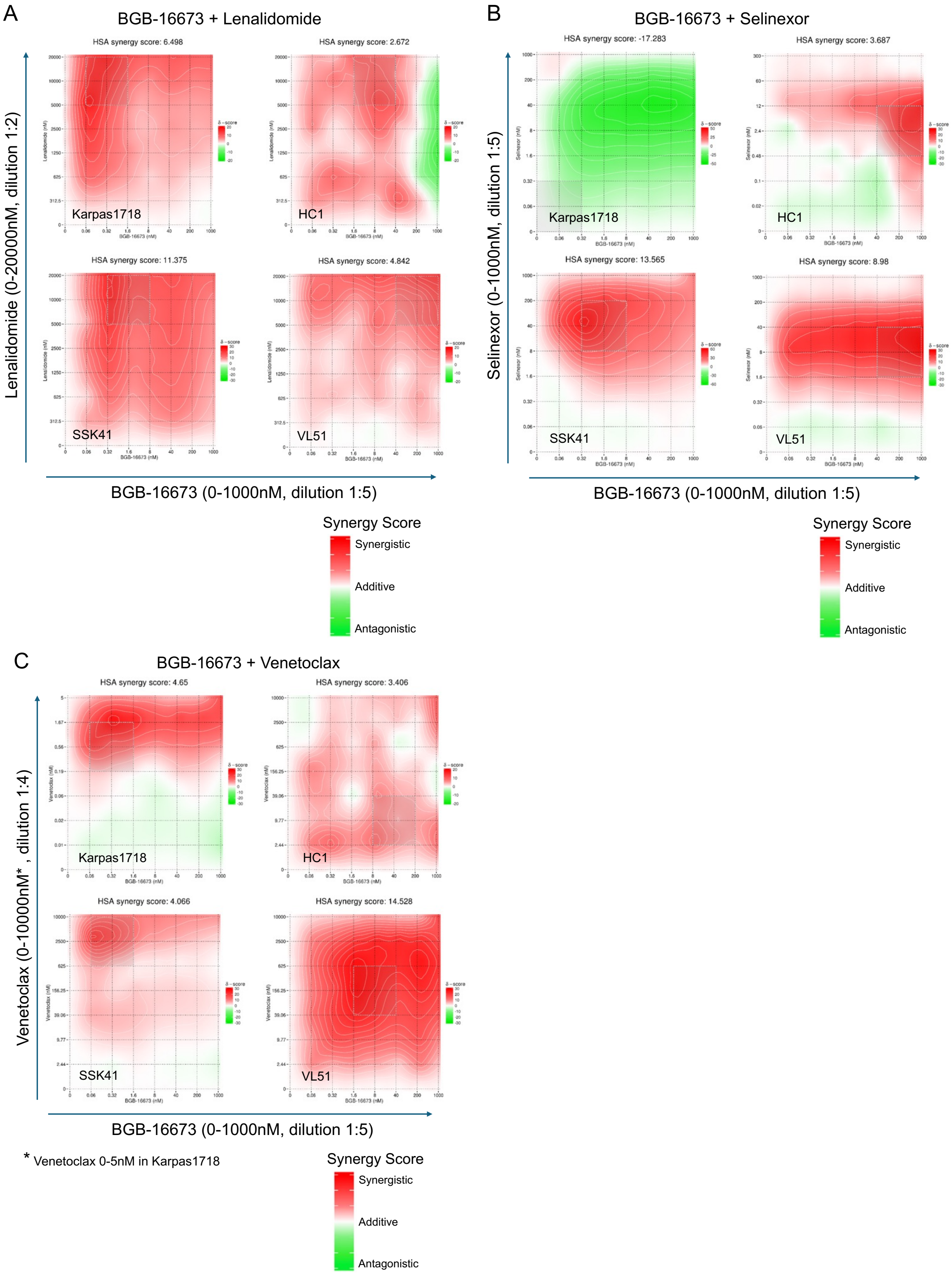

D

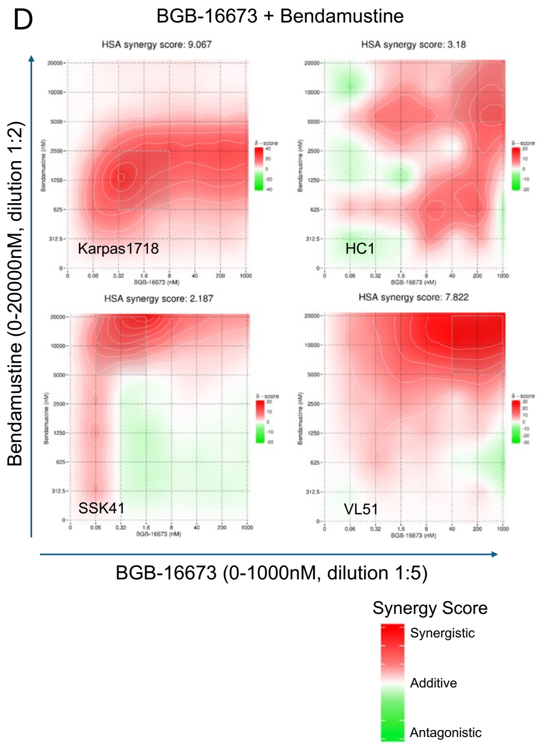
BGB-16673 + Bendamustine
HSA synergy score: 9.067
Karpas1718
HSA synergy score: 3.18
HC1
HSA synergy score: 2.187
SSK41
HSA synergy score: 7.822
VL51
Bendamustine (0-20000nM, dilution 1:2)
BGB-16673 (0-1000nM, dilution 1:5)
Synergy Score
Synergistic
Additive
Antagonistic

E

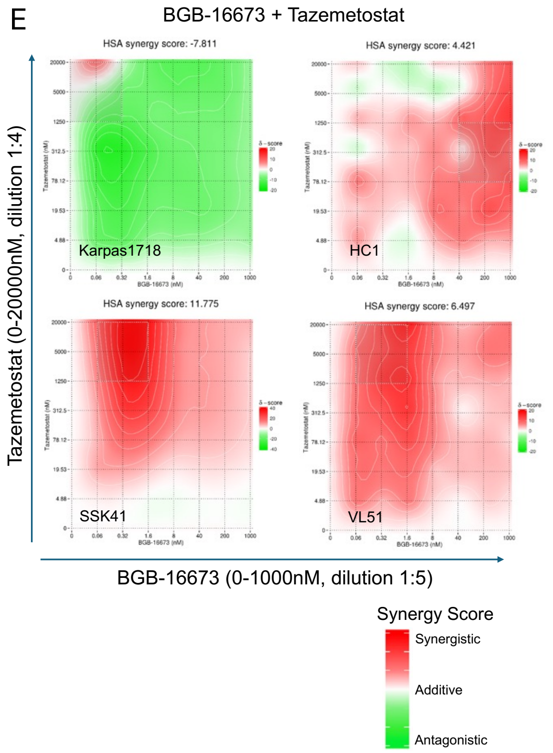
BGB-16673 + Tazemetostat
HSA synergy score: -7.811
Karpas1718
HSA synergy score: 4.421
HC1
HSA synergy score: 11.775
SSK41
HSA synergy score: 6.497
VL51
Tazemetostat (0-20000nM, dilution 1:4)
BGB-16673 (0-1000nM, dilution 1:5)
Synergy Score
Synergistic
Additive
Antagonistic

F

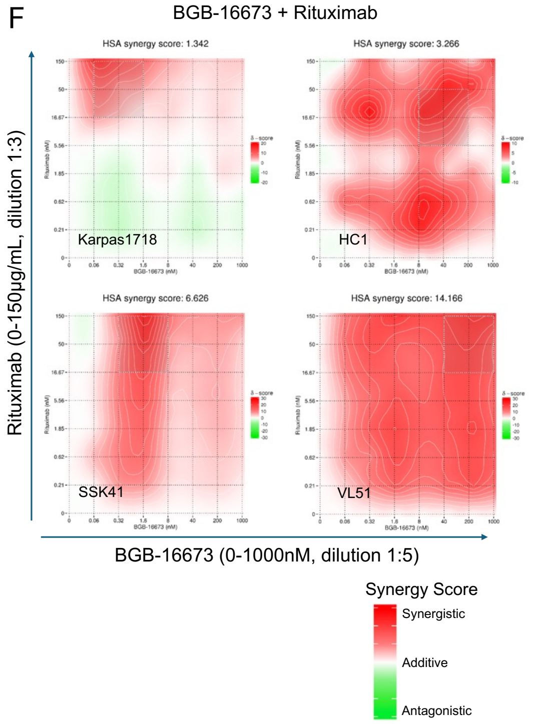
BGB-16673 + Rituximab
HSA synergy score: 1.342
Karpas1718
HSA synergy score: 3.266
HC1
HSA synergy score: 6.626
SSK41
HSA synergy score: 14.166
VL51
Rituximab (0-150μg/mL, dilution 1:3)
BGB-16673 (0-1000nM, dilution 1:5)
Synergy Score
Synergistic
Additive
Antagonistic

**Supplementary Figure 7.** Apoptosis induction by BGB-16673 as a single agent and in combination with venetoclax, sonrotoclax, or bendamustine in Karpas1718 (upper panel) and in the VL51 (lower panel) cell lines. Barplots show the mean of two biological replicates after 24 hours of treatment. Error bars represent the standard deviation of the mean.

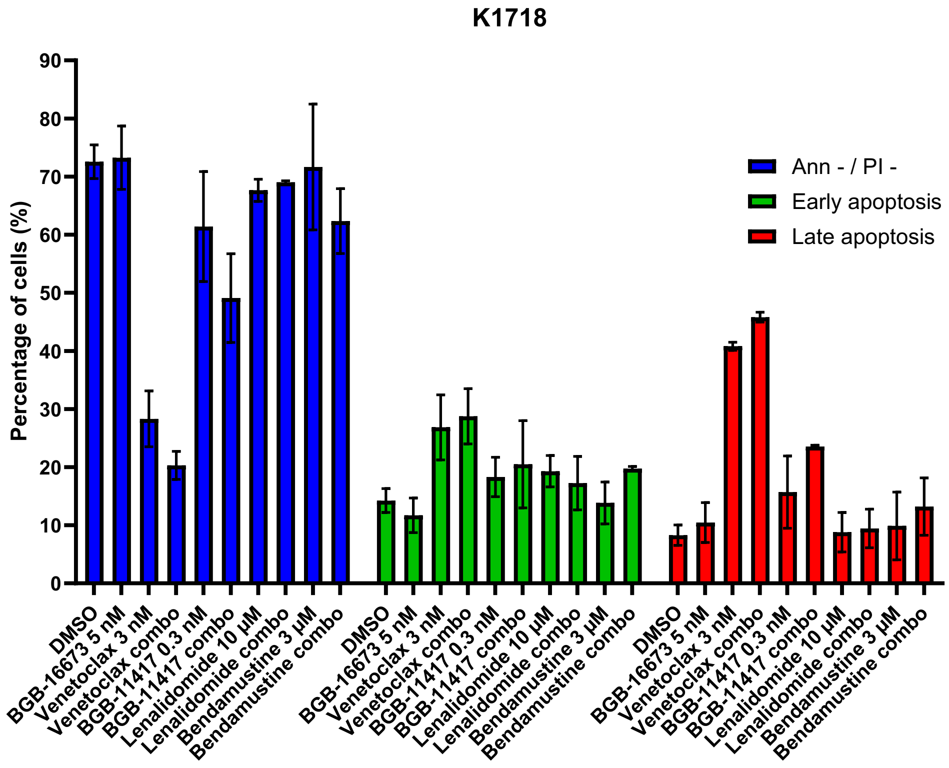


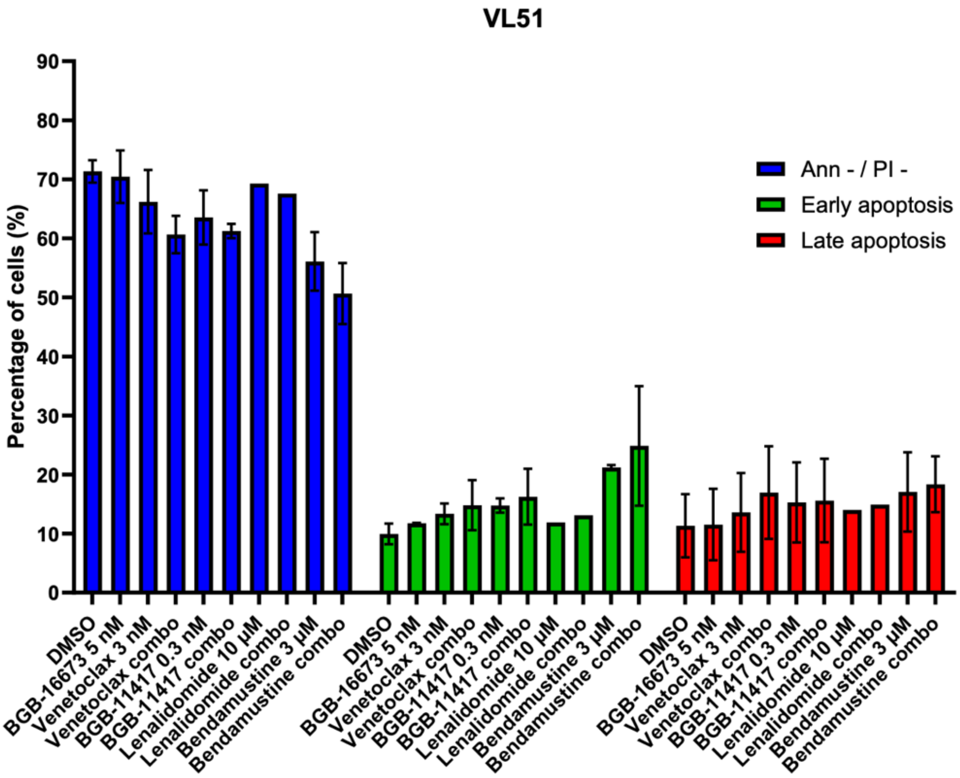

**Supplementary Figure 8.** Effect of the BTK-degrader BGB-16673 and the BCL2 inhibitor Sonrotoclax on the protein levels of BTK and pBTK in the two MZL cell lines Karpas1718 (24hr, A) and VL51 (24hr, B). Immunoblotting is representative of two independent experiments. The bar plot at the bottom shows the mean of the protein quantification in the two experiments. Error bars correspond to the standard deviation of the mean, and numbers represent the p-value from a t-test comparing each treatment to the control (DMSO, grey bar).

A) KARPAS1718

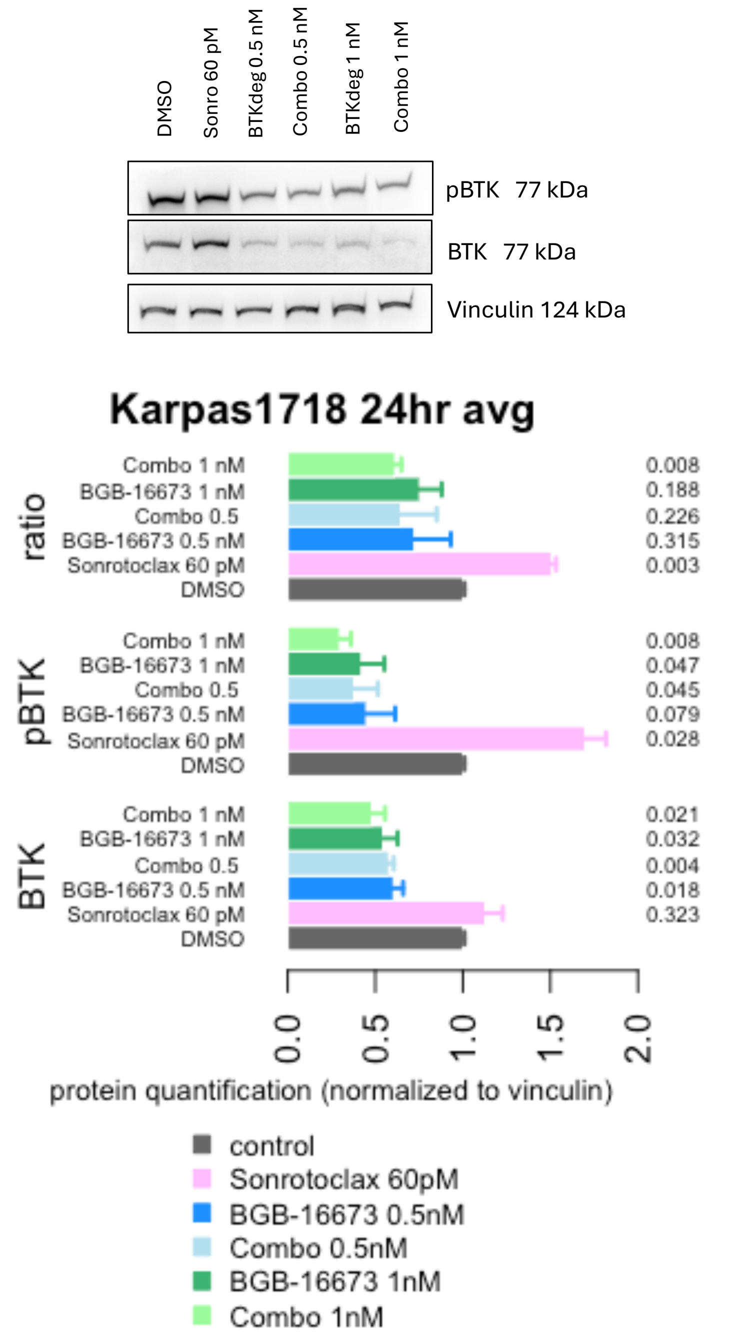

B)

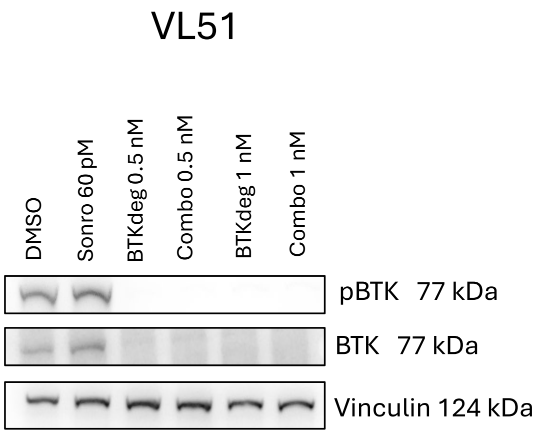

VL51
DMSO
Sonro 60 pM
BTKdeg 0.5 nM
Combo 0.5 nM
BTKdeg 1 nM
Combo 1 nM
pBTK 77 kDa
BTK 77 kDa
Vinculin 124 kDa


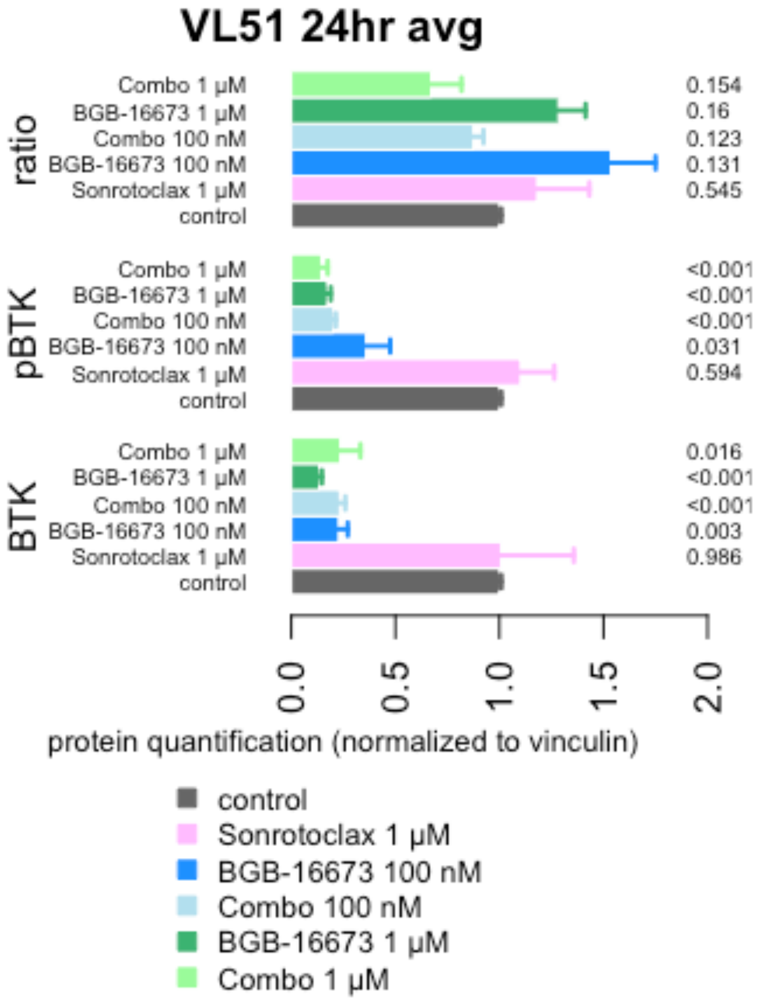

VL51 24hr avg
ratio
Combo 1 μM
BGB-16673 1 μM
Combo 100 nM
BGB-16673 100 nM
Sonrotoclax 1 μM
control
0.154
0.16
0.123
0.131
0.545
pBTK
Combo 1 μM
BGB-16673 1 μM
Combo 100 nM
BGB-16673 100 nM
Sonrotoclax 1 μM
control
<0.001
<0.001
<0.001
0.031
0.594
BTK
Combo 1 μM
BGB-16673 1 μM
Combo 100 nM
BGB-16673 100 nM
Sonrotoclax 1 μM
control
0.016
<0.001
<0.001
0.003
0.986
0.0
0.5
1.0
1.5
2.0
protein quantification (normalized to vinculin)
control
Sonrotoclax 1 μM
BGB-16673 100 nM
Combo 100 nM
BGB-16673 1 μM
Combo 1 μM

**Supplementary Figure 9.** Effect of the BTK-degrader BGB-16673 and the BCL2 inhibitor sonrotoclax on the protein levels of BCL2, MCL1 and BCLXL in the two MZL cell lines Karpas1718 (24hr, A) and VL51 (24hr, B). Immunoblotting is representative of two independent experiments. The bar plot at the bottom shows the mean of the protein quantification in the two experiments. Error bars correspond to the standard deviation of the mean, and numbers represent the p-value from a t-test comparing each treatment to the control (DMSO, grey bar).

A)

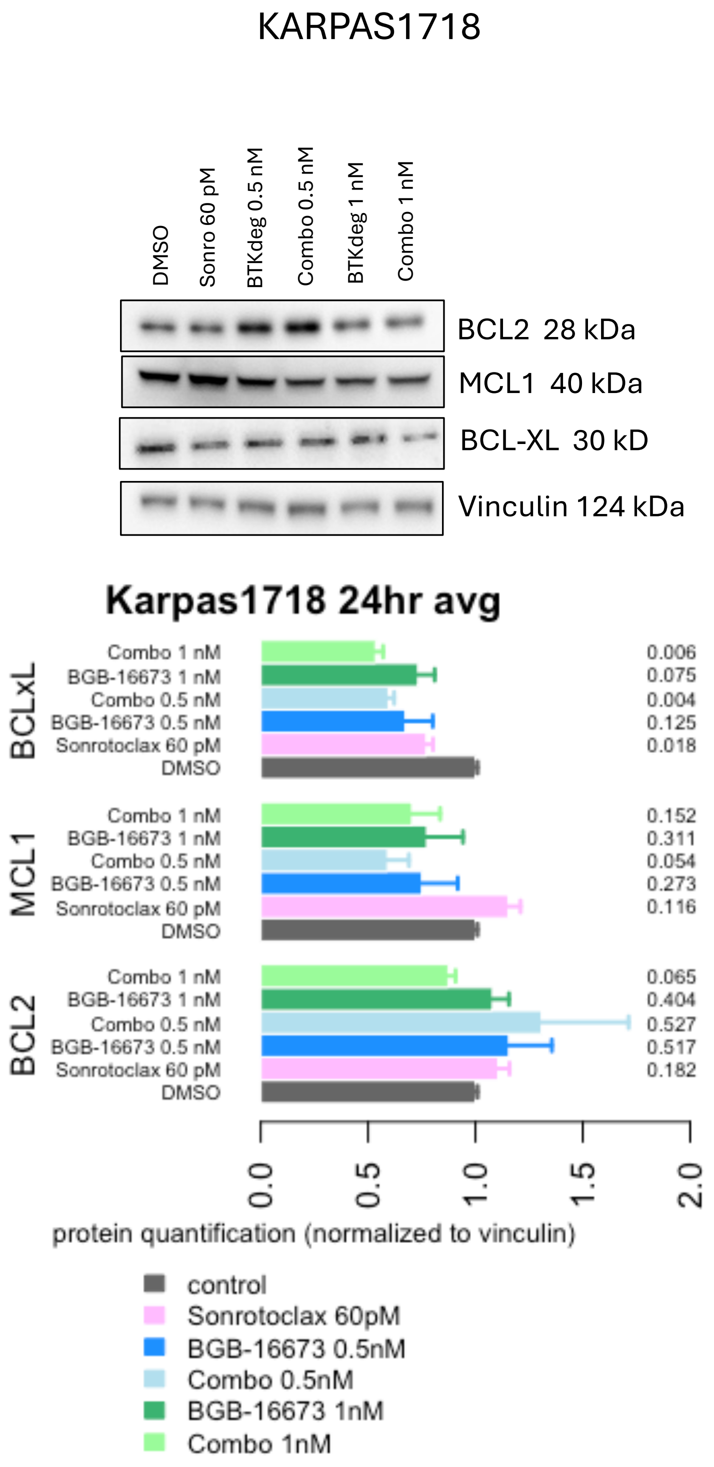

B)

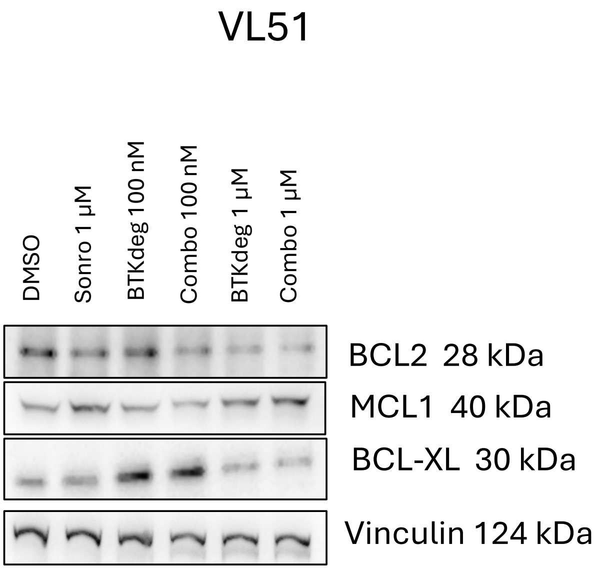

VL51
DMSO
Sonro 1 μM
BTKdeg 100 nM
Combo 100 nM
BTKdeg 1 μM
Combo 1 μM
BCL2 28 kDa
MCL1 40 kDa
BCL-XL 30 kDa
Vinculin 124 kDa


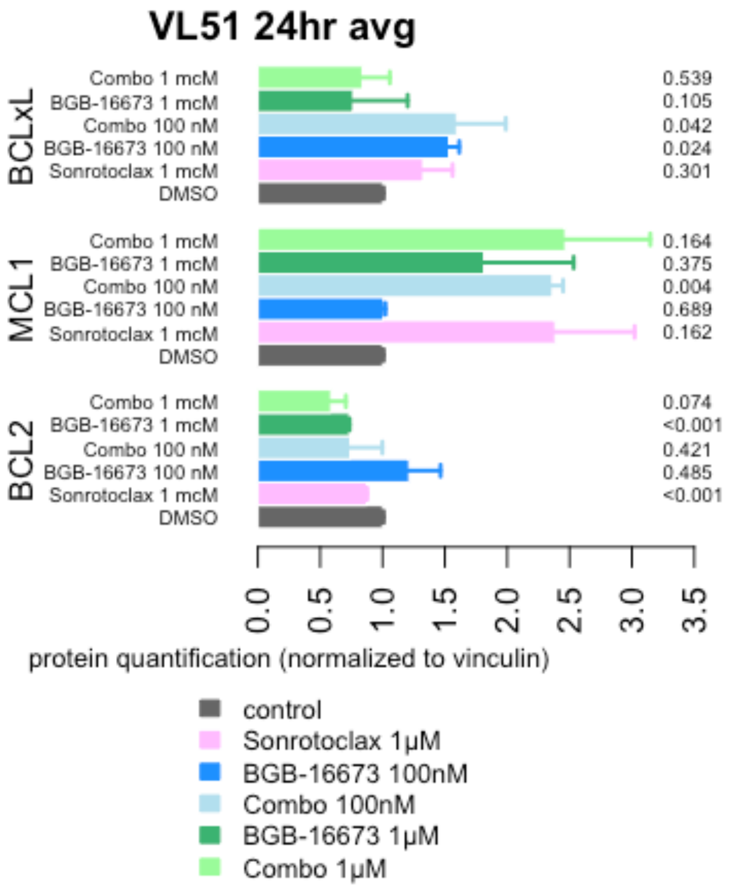

VL51 24hr avg
BCLxL
Combo 1 mcM
BGB-16673 1 mcM
Combo 100 nM
BGB-16673 100 nM
Sonrotoclax 1 mcM
DMSO
0.539
0.105
0.042
0.024
0.301
MCL1
Combo 1 mcM
BGB-16673 1 mcM
Combo 100 nM
BGB-16673 100 nM
Sonrotoclax 1 mcM
DMSO
0.164
0.375
0.004
0.689
0.162
BCL2
Combo 1 mcM
BGB-16673 1 mcM
Combo 100 nM
BGB-16673 100 nM
Sonrotoclax 1 mcM
DMSO
0.074
<0.001
0.421
0.485
<0.001
0.0
0.5
1.0
1.5
2.0
2.5
3.0
3.5
protein quantification (normalized to vinculin)
control
Sonrotoclax 1μM
BGB-16673 100nM
Combo 100nM
BGB-16673 1μM
Combo 1μM

**Supplementary Figure 10.** Superimposition of the binding pose of BGB-16673 predicted by docking calculations with the crystallographic binding mode of L0Z (PDB ID: 6S90) to BTK. The BTK protein is depicted as light blue ribbons, with key residues highlighted in blue sticks. BGB-16673 and L0Z are represented by purple and orange sticks, respectively. Key receptor–ligand interactions are depicted as black dashed lines. For clarity, only hydrogen atoms directly involved in these interactions are shown. The protein-drug interaction sites are highlighted as insets.

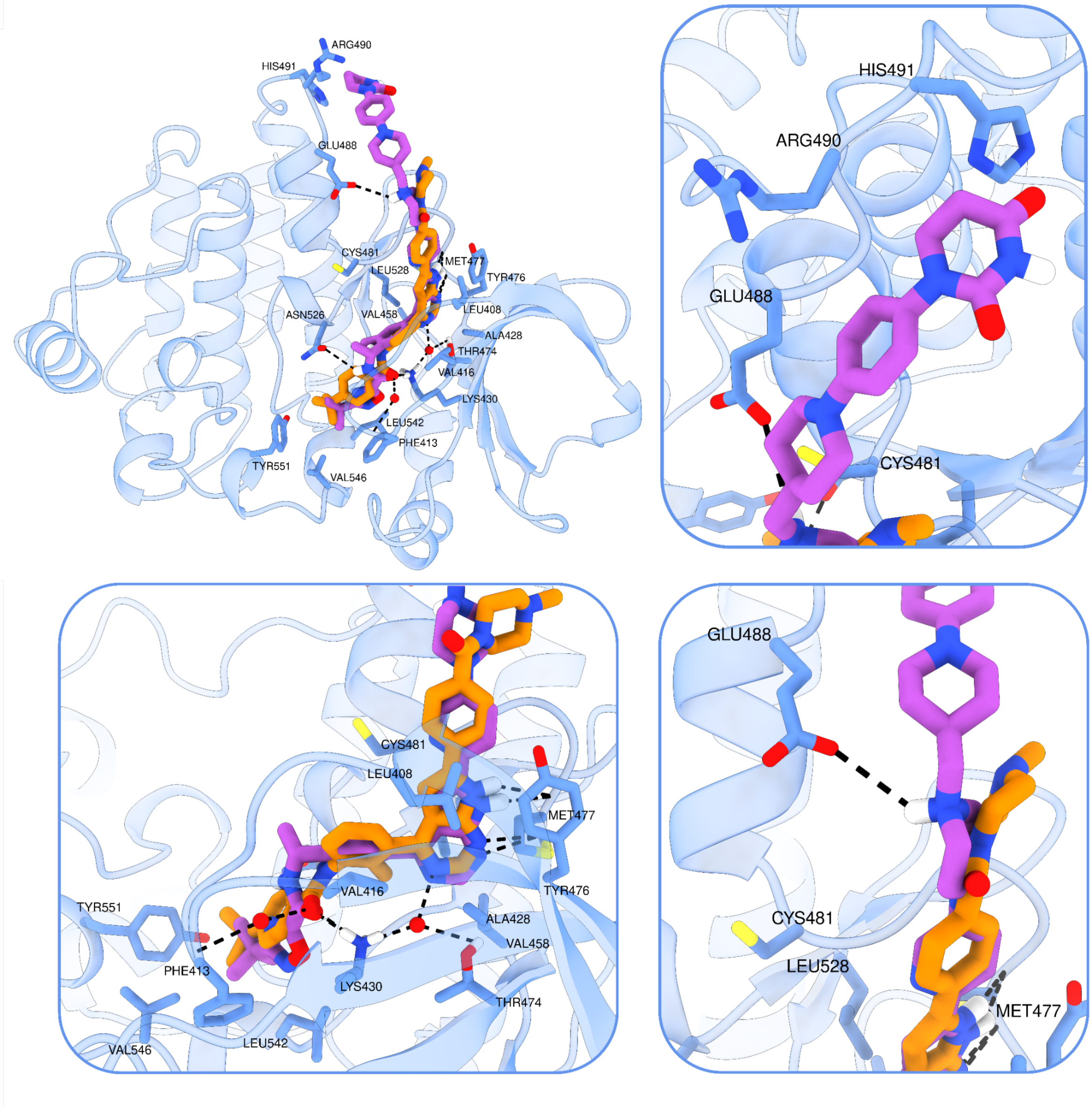

**Supplementary Table S1. Drug concentrations range and dilutions used in the single treatments and in the combinations with the BTK-degrader BGB-16673.**

| experiment | compound | cell line | max concentration | drug diluton |
|---|---|---|---|---|
| single | BGB-16673 | all models | 10000nM | dil 1/10 |
| single | BGB-21704 | Karpas1718, SSK4 | 10000nM | dil 1/10 |
| single | Zanubrutinib | Karpas1718, SSK4 | 10000nM | dil 1/10 |
| single | Sonrotoclax | Karpas1718, SSK4 | 10000 (5nM in Karpas1718) | dil 1/4 (dil 1/3 in Karpas1718) |
| single | Venetoclax | Karpas1718, SSK4 | 10000 (5nM in Karpas1718) | dil 1/4 (dil 1/3 in Karpas1718) |
| combo with BGB-16673 (0-1000nM dil 1/5) | Lenalidomide | Karpas1718, SSK4 | 20000nM | dil 1/2 |
| combo with BGB-16673 (0-1000nM dil 1/5) | Selinexor | Karpas1718, SSK4 | 1000nM (300nM in HC1) | dil 1/5 |
| combo with BGB-16673 (0-1000nM dil 1/5) | Venetoclax | Karpas1718, SSK4 | 10000 (5nM in Karpas1718) | dil 1/4 (dil 1/3 in Karpas1718) |
| combo with BGB-16673 (0-1000nM dil 1/5) | Sonrotoclax | Karpas1718, SSK4 | 10000 (5nM in Karpas1718) | dil 1/4 (dil 1/3 in Karpas1718) |
| combo with BGB-16673 (0-1000nM dil 1/5) | Bendamustine | Karpas1718, SSK4 | 20000nM | dil 1/2 |
| combo with BGB-16673 (0-1000nM dil 1/5) | Tazemetostat | Karpas1718, SSK4 | 20000nM | dil 1/4 (dil 1/3 in Karpas1718) |
| combo with BGB-16673 (0-1000nM dil 1/5) | Rituximab | Karpas1718, SSK4 | 150µg/mL | dil 1/3 |

| experiment | compound | cell line | max concentration | drug diluton |
|---|---|---|---|---|
| **single** | BGB-16673 | all models | 10000nM | dil 1/10 |
| **single** | BGB-21704 | Karpas1718, SSK41 | 10000nM | dil 1/10 |
| **single** | Zanubrutinib | Karpas1718, SSK41 | 10000nM | dil 1/10 |
| **single** | Sonrotoclax | Karpas1718, SSK41, HC1, VL51 | 10000 (5nM in Karpas1718) | dil 1/4 (dil 1/3 in Karpas1718) |
| **single** | Venetoclax | Karpas1718, SSK41, HC1, VL51 | 10000 (5nM in Karpas1718) | dil 1/4 (dil 1/3 in Karpas1718) |
| **combo with BGB-16673 (0-1000nM dil 1/5)** | Lenalidomide | Karpas1718, SSK41, HC1, VL51 | 20000nM | dil 1/2 |
| **combo with BGB-16673 (0-1000nM dil 1/5)** | Selinexor | Karpas1718, SSK41, HC1, VL51 | 1000nM (300nM in HC1) | dil 1/5 |
| **combo with BGB-16673 (0-1000nM dil 1/5)** | Venetoclax | Karpas1718, SSK41, HC1, VL51 | 10000 (5nM in Karpas1718) | dil 1/4 (dil 1/3 in Karpas1718) |
| **combo with BGB-16673 (0-1000nM dil 1/5)** | Sonrotoclax | Karpas1718, SSK41, HC1, VL51 | 10000 (5nM in Karpas1718) | dil 1/4 (dil 1/3 in Karpas1718) |
| **combo with BGB-16673 (0-1000nM dil 1/5)** | Bendamustine | Karpas1718, SSK41, HC1, VL51 | 20000nM | dil 1/2 |
| **combo with BGB-16673 (0-1000nM dil 1/5)** | Tazemetostat | Karpas1718, SSK41, HC1, VL51 | 20000nM | dil 1/4 (dil 1/3 in Karpas1718) |
| **combo with BGB-16673 (0-1000nM dil 1/5)** | Rituximab | Karpas1718, SSK41, HC1, VL51 | 150µg/mL | dil 1/3 |

**Supplementary Table S2. IC50 values obtained after five days of exposure to the BTK-degrader BGB-16673 or to the BTK inhibitor Zanubrutinib in cell lines derived from MZL (VL51, SSK41, Karpas1718, HC1, HAIRM, ESKOL) and from MCL (REC1, MINO).**

| compound | sample | IC50 (nM) | se (nM) |
|---|---|---|---|
| BGB-16673 | Karpas1718 | 1.8866 | 0.1505 |
| | SSK41 | 0.2668 | 0.1015 |
| | VL51 | >1000 | - |
| | ESKOL | >1000 | - |
| | HAIRM | >1000 | - |
| | HC1 | >1000 | - |
| | REC1 | 0.0014 | 0.0170 |
| | MINO | 0.0315 | 0.0075 |
| Zanubrutinib | Karpas1718 | 2.5486 | 2.5486 |
| | SSK41 | 0.1592 | 0.1592 |
| | REC1 | 0.2204 | 0.2204 |
| | MINO | 2.2631 | 2.2631 |

**Supplementary Table S3 (Excel file).** Limma analysis and Gene Set Enrichment Analyses (GSEA) of RNA-Seq data from the MZL cell line Karpas1718 exposed to BGB-16673 (5 nM) or zanubrutinib (5 nM) compared to DMSO. Cells were treated for 8, 12, and 24 hours.

**Supplementary Table S4. IC50 values obtained after five days of exposure to the BCL2 inhibitors sonrotoclax and venetoclax in six MZL cell lines.**

| compound | histology | cell line | IC50 (nM) | se (nM) |
|---|---|---|---|---|
| Sonrotoclax | MZL | VL51 | 1646.4 | 431.9 |
| | MZL | Karpas1718 | 0.0218 | 0.0233 |
| | MZL | SSK41 | 6281.6 | 1142.2 |
| | MZL | HC1 | 1.6540 | 1.8719 |
| | MZL | HAIRM | 0.0195 | 0.0202 |
| | MZL | ESKOL | 16015.1 | 4437.8 |
| Venetoclax | MZL | VL51 | 2446.3 | 291.7 |
| | MZL | Karpas1718 | 1.0663 | 0.2936 |
| | MZL | SSK41 | 5494.1 | 1025.5 |
| | MZL | HC1 | 1129.7 | 491.2 |
| | MZL | HAIRM | 20.085 | 13.028 |
| | MZL | ESKOL | 17158.4 | 4536.3 |
